# Non-Neural Sources Systematically Impact Aperiodic EEG Activity

**DOI:** 10.64898/2026.01.29.702285

**Authors:** Marius Tröndle, Nicolas Langer

**Author notes:** CORRESPONDING AUTHOR: Marius Tröndle, Methods of Plasticity Research, Department of Psychology, University of Zurich, Binzmühlenstrasse 14, CH-8050, Zürich, Switzerland.

## Abstract

Aperiodic, 1/f-like EEG activity has emerged as a key index of neural population dynamics, offering insights into excitation/inhibition balance, cognitive states, and clinical conditions. However, the foundational assumption that these parameters reflect neural dynamics rather than non-neural sources remains largely untested. Here, we systematically quantify how data quality and physiological artifacts influence aperiodic parameter estimation across two independent datasets (N=99 and N=103). Findings converged across data sets and two complementary analytical approaches: experimental artifact manipulation and selective signal decomposition.

We demonstrate that ocular artifacts and poor data quality systematically inflate aperiodic offsets and exponents, while muscular artifacts exert the opposite effects. These spatially widespread influences persist after state-of-the-art preprocessing and reach magnitudes comparable to reported neurophysiologically meaningful group differences.

Our results reveal that differential rates of non-neural sources can drive spurious neural interpretations; consequently, we introduce and validate an approach to mitigate these biases, guiding valid inference in basic and clinical aperiodic research.

## Introduction

Electroencephalography (EEG) offers a window into multi-scale dynamics of the human brain, traditionally characterized by periodic oscillations. However, a transformative shift in neurophysiology has revealed that the “background” aperiodic signal—defined by its 1/f-like power-law scaling—is not stochastic noise, but a biologically rich index of neural population dynamics^1,2^. This aperiodic component is now recognized as a critical proxy for the global balance of excitation and inhibition (E/I), providing a unique physiological signature of cognitive states and a wide array of clinical conditions. As computational methods for isolating these components have matured, aperiodic activity has moved from the periphery to the center of neuroscientific inquiry.

These aperiodic signals are typically parameterized by two components: the offset, reflecting broadband power shifts, and the exponent, describing spectral steepness. While the offset has been linked to neuronal population spiking rates^3^, variations in the exponent are proposed to reflect differences in neural noise^4,5^, self-organized criticality^6–8^, or the most prominently^9^, the balance between inhibitory and excitatory synaptic activity^10,11^. Empirically, these parameters have demonstrated remarkable sensitivity to aging^12–14^, task-related neural dynamics^15–17^, and diverse clinical conditions, showing alterations in autism spectrum disorders (ASD)^18,19^, schizophrenia^20,21^, attention-deficit/hyperactivity disorder (ADHD)^22,23^, and neurodegenerative diseases^24–26^. This has positioned aperiodic activity as a high-potential tool for diagnostic and treatment monitoring. The potential for these metrics to serve as reliable biomarkers, however, hinges on the assumption that they provide a high-fidelity reflection of neural processes.

Yet, EEG recordings are inherently susceptible to non-neural sources, including ocular, muscular, and cardiac artifacts, as well as measurement noise^27^, all of which can distort spectral estimates. Initial evidence suggests that these non-neural sources indeed extend to aperiodic estimates; for instance, recent machine learning models can discriminate between high- and low-quality data segments based on aperiodic features^28^, indicating an inherent sensitivity to data quality. However, while these findings signal potential contamination, only isolated studies have directly implicated specific sources, such as cardiac artifacts^29^, leaving a comprehensive assessment lacking.

Consequently, it remains unknown how specific artifact sources and variations in data quality systematically affect aperiodic parameter estimates. This knowledge gap represents a critical vulnerability in the field: given the well-documented differences in data quality or artifact prevalence between patient and control groups^30,31^, or even between experimental conditions, failing to quantify these influences may lead to spurious neurobiological interpretations of what are, in fact, non-neural differences. Many of the empirical patterns currently cited as evidence for shifts in excitation/inhibition (E/I) balance^9,17,32^ may instead reflect systematic differences in physiological artifacts or recording quality. Disentangling these influences is therefore a prerequisite for advancing our fundamental understanding of aperiodic parameters To address this challenge, we systematically quantified how data quality and physiological artifacts influence aperiodic parameter estimation in resting-state EEG across two large independent datasets. As illustrated in Figure 1, we first assessed the impact of data quality metrics on aperiodic parameters. Second, we examined how ocular, muscular, and cardiac activity artifacts alter aperiodic estimates. We implemented two complementary methodological frameworks: a *component retention* approach, selectively removing or retaining independent components (ICs) linked to specific artifacts, and an *incremental contamination* approach, that experimentally manipulated contamination levels by systematically mixing artifact-containing segments into otherwise clean data at predefined levels.

**Figure 1:**
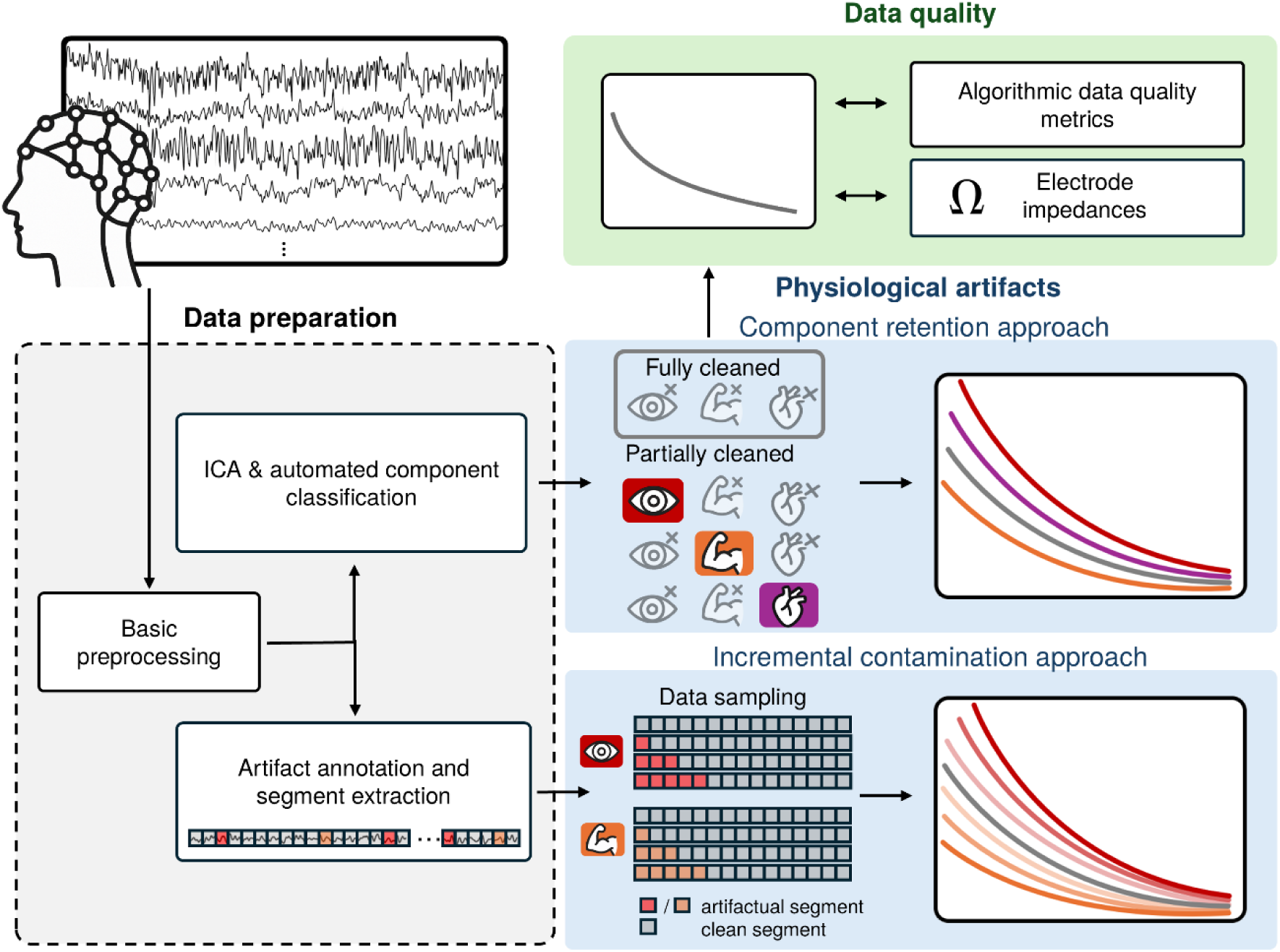
Analytical Framework for quantifying non-neural influences on aperiodic parameters. Schematic overview of the two complementary methodological approaches assessing the impact of data quality (green box) and specific physiological artifacts (blue boxes) on aperiodic parameters estimation. Basic preprocessing (gray box) included 0.5 Hz high pass filtering and removal of bad channels and of line noise. For data quality analyses, the aperiodic parameters were derived from fully cleaned EEG data (all artifactual ICs removed; green box, left). In the component retention approach (blue top), specific aperiodic parameters were estimated from signals where target artifact ICs were selectively retained (colored symbols) or removed (crossed and faded symbols). In the incremental contamination approach (blue bottom), artifact-containing segments were systematically titrated into clean EEG data at four predefined levels (0, 1, 3, or 5 artifact segments out of 15 total), prior parameter estimation. Note that cardiac artifacts were excluded from the incremental contamination approach due to specific signal characteristics (see Methods).

Together, these analyses provide the first comprehensive quantification of how specific artifact sources and variations in data quality shape aperiodic parameter estimates. Our results clarify the non-neural sources of aperiodic activity and introduce and validate an approach to mitigate these biases, providing practical methodological guidance for robust inference in basic and clinical research.

## Results

### Final sample characteristics

To ensure the robustness and generalizability of our findings, we analyzed two independent EEG datasets (N = 99 and N = 103) recorded with distinct hardware configurations. The main dataset utilized an electrolyte gel-based EEG system and included concurrent eye-tracking, electromyography (EMG), and electrocardiography (ECG) to provide high-fidelity ground truth for physiological artifact identification. The validation dataset utilized a saline-based EEG system and included comparable EEG and eye-tracking data but lacked EMG and ECG recordings.

Following the exclusion of low-quality recordings at the end of preprocessing, the final samples consisted of 89 participants in the main dataset (age range 18.1 to 38.7 years, sd=3.8, 67 female), and 93 participants in the validation dataset (age range 19.7 to 32.2 years, sd = 3.0, 57 female). Sample sizes for specific sub-analysis varied based on the availability of electrode impedance measures and the presence of detectable artifacts (see Supplementary Table 1 for a detailed breakdown).

Crucially, our findings were highly consistent across both datasets. For the sake of conciseness, statistical results are reported for both samples throughout, while visualizations in the main text focus on the main dataset due to its comprehensive physiological coverage and superior artifact detection capabilities. Full cross-dataset visualizations are provided in Supplement 3.

### Effect of data quality metrics on the aperiodic signal component

We first examined the relationship between data quality and the aperiodic parameter estimates using both a hardware-based measure (electrode impedances) and three algorithmic quality metrics: proportion of overall high amplitude datapoints (OHA), proportion of timepoints of high variance (THV), and proportion of channels of high variance (CHV). Importantly, all aperiodic parameters and algorithmic quality metrics were estimated from fully cleaned data (i.e. after removal of all ICs identified as artifacts by ICLabel^33^) to assess whether the effects of data quality persisted even after applying state-of-the-art preprocessing methods.

#### Hardware based quality metrics: Electrode impedances

Electrode impedances were not significantly related to the aperiodic exponent in either dataset. However, significant positive associations were found with the aperiodic offset in the main dataset, with a similar trend in the validation dataset (see Table 1). As shown in the topographical maps (Figure 2B), this association was broadly distributed across the scalp.

**Figure 2:**
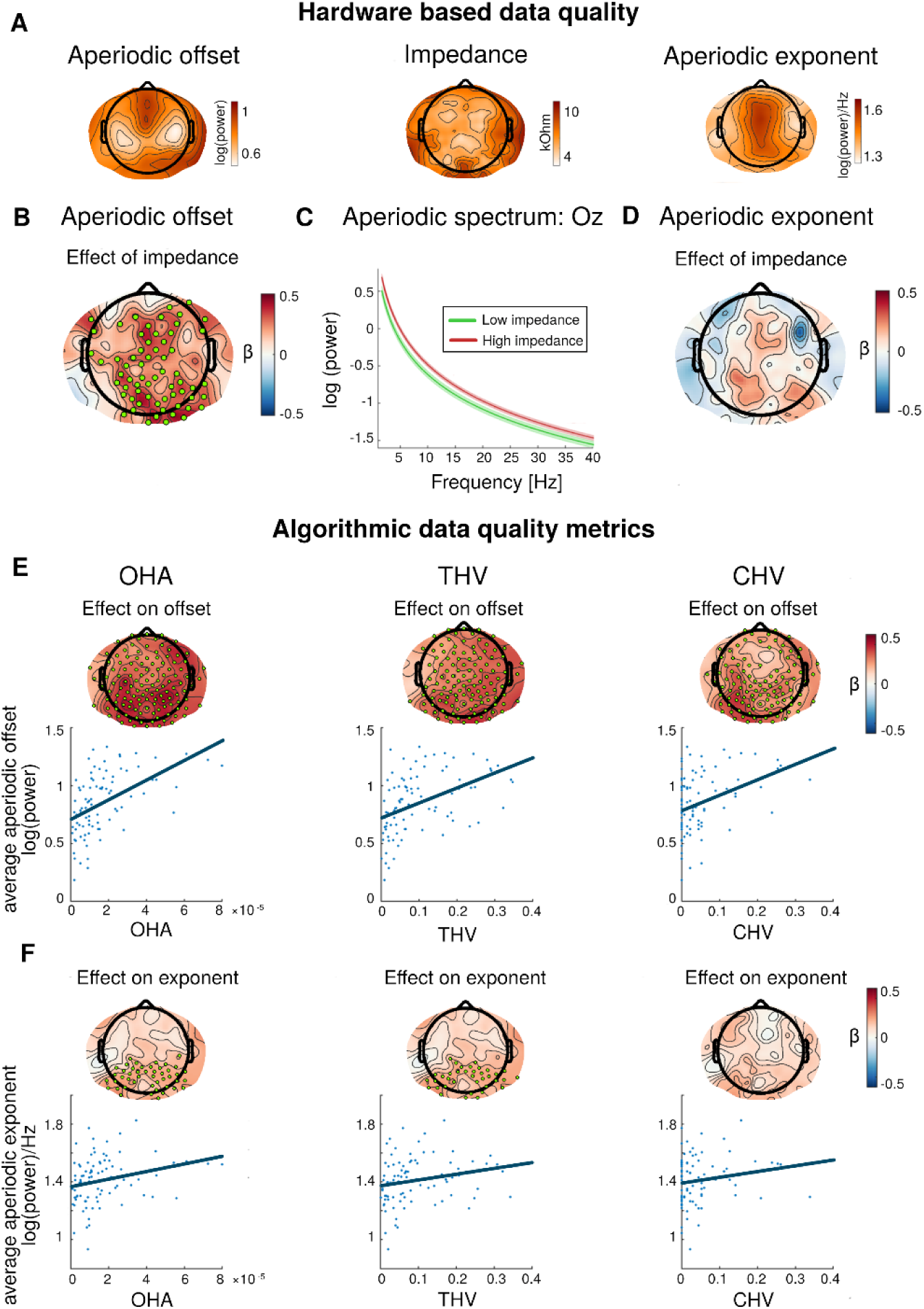
Impact of data quality on aperiodic parameter estimation. All visualizations represent findings from the main dataset. Panels A-D illustrate the relationship between electrode impedances and the aperiodic parameters. Panel A shows the topographical distribution of the aperiodic offset, exponent, and electrode impedances. Panels B&D display standardized regression coefficients (β) predicting the aperiodic parameters from electrode impedances; green dots denote significant electrode clusters identified via cluster-based permutation tests (CBPT). Panel C presents median-split aperiodic spectra at electrode Oz, shaded areas represent standard error of the mean. Panels E-F depict the influence of algorithmic quality metrics (OHA THV, CHV) on aperiodic parameters. Topographies show standardized β coefficients, with green dots indicating significant electrodes of the CBPTs. Scatterplots illustrate the relationship between quality metrics and aperiodic parameters averaged across all electrodes.

**Table 1:** Statistical results (Cluster based permutation tests, CBPTs): Effect of data quality metrics on the aperiodic signal parameters.

| | Aperiodic offset<br>(avg. standardized $\beta$, cluster p) | | Aperiodic exponent<br>(avg. standardized $\beta$, cluster p) | |
| --- | --- | --- | --- | --- |
|  | Main dataset | Validation dataset | Main dataset | Validation dataset |
| Electrode impedance | $\bar{\beta} = 0.31$,<br>$p \leq 0.001$ | $\bar{\beta} = 0.40$,<br>$p = 0.100$ | $\bar{\beta} = 0.26$,<br>$p = 0.115$ | $\bar{\beta} = -0.33$,<br>$p = 0.250$ |
| OHA | $\bar{\beta} = 0.40$,<br>$p \leq 0.001$ | $\bar{\beta} = 0.48$,<br>$p \leq 0.001$ | $\bar{\beta} = 0.29$,<br>$p = 0.009$ | $\bar{\beta} = 0.30$,<br>$p \leq 0.001$ |
| THV | $\bar{\beta} = 0.35$,<br>$p \leq 0.001$ | $\bar{\beta} = 0.51$,<br>$p \leq 0.001$ | $\bar{\beta} = 0.24$,<br>$p = 0.003$ | $\bar{\beta} = 0.29$,<br>$p = 0.003$ |
| CHV | $\bar{\beta} = 0.30$,<br>$p \leq 0.001$ | $\bar{\beta} = 0.34$,<br>$p \leq 0.001$ | $\bar{\beta} = 0.23$,<br>$p = 0.100$ | $\bar{\beta} = 0.25$,<br>$p = 0.028$ |

*Note: Standardized $\beta$ 's are averaged across all electrodes of the significant cluster. If no cluster was significant, $\beta$ 's are averaged across the largest non-significant cluster.*

#### Algorithmic quality metrics

Higher rates of OHA, CHV and THV, indicating poorer data quality, were significantly associated with higher aperiodic offsets and steeper aperiodic exponents in both datasets (Table 1, Figure 2E-F). These influences were spatially widespread, though notably most pronounced over parietal and occipital regions. This confirms that variations in data quality systematically bias the estimated steepness of the aperiodic component even after rigorous cleaning.

### Effect of physiological artifacts on the aperiodic signal component

We next examined how physiological artifacts influence the aperiodic parameter estimation using two complementary approaches illustrated in Figure 1. Both the component retention and incremental contamination methods yielded highly consistent results across datasets, revealing substantial and systematic biases.

#### Ocular artifacts

In the component retention approach, ocular contamination significantly affected both the aperiodic offset and the aperiodic exponent (Table 2). Ocular artifacts were associated with larger offsets and steeper aperiodic slopes. Crucially, these effects were not restricted to prefrontal electrodes near the eyes, but extended across the scalp to encompass temporal, parietal, and occipital regions (Figure 3B and 3D). This indicates that ocular activity distorts the aperiodic signal globally, even during eyes-closed resting-state recordings. These effects were predictably amplified in the eyes-open data (Supplement 4).

**Figure 3:**
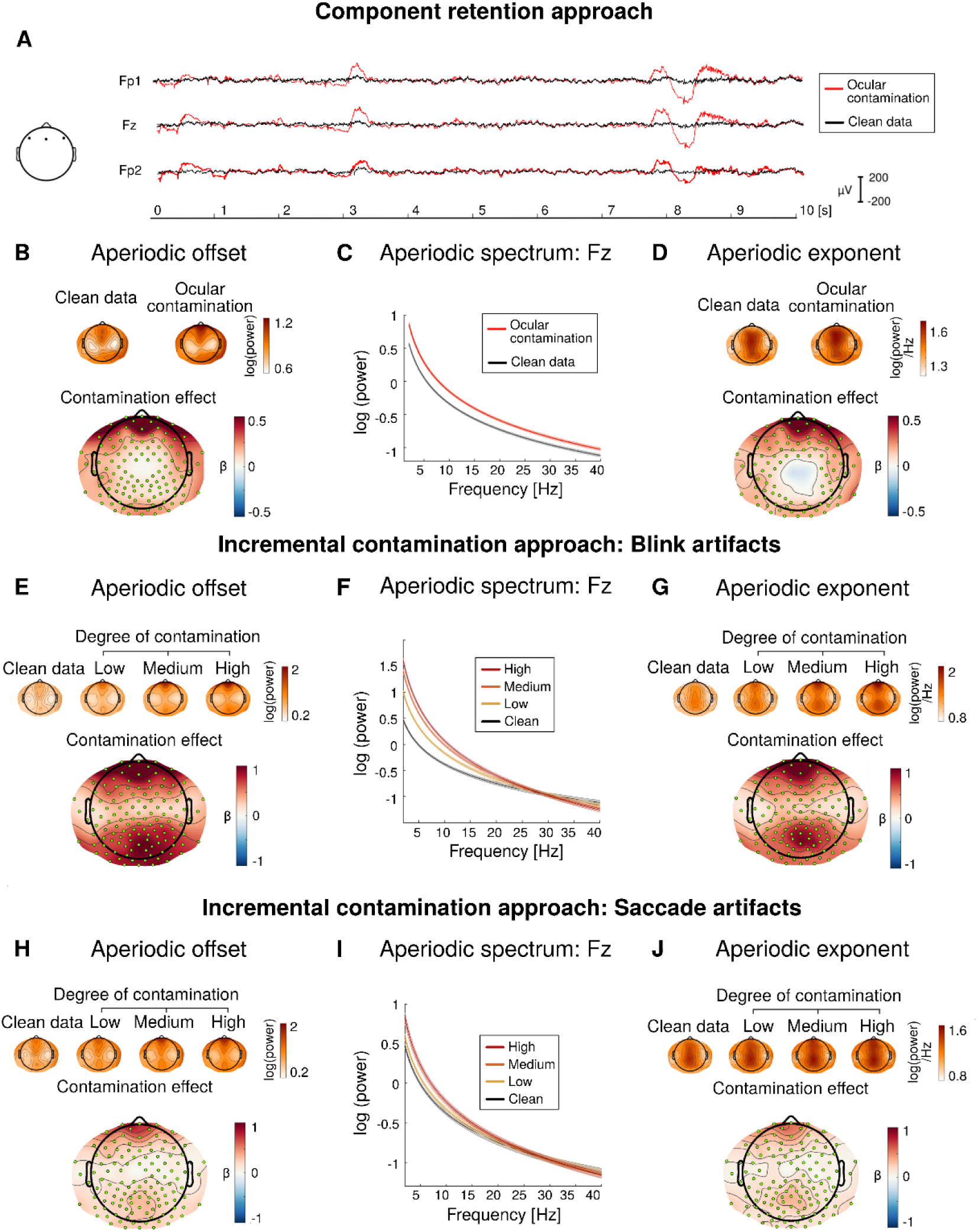
Impact of ocular artifact contamination on aperiodic parameter estimation. Visualizations are based on the main dataset. Panels A-D illustrate findings from the component retention approach. Panel A shows data from three prefrontal electrodes (indicated on head schematic from left to right: Fp1, Fz, Fp2) before (red) and after (black) removal of ICs classified as ocular activity. Small topographies on top of B&D display aperiodic parameters derived from fully cleaned data (all artifactual ICs removed) and partially cleaned data (ocular ICs retained). Large maps display standardized regression coefficients from linear mixed-effects models predicting aperiodic parameters from ocular contamination; green dots mark electrodes significant in CBPT. Panel C shows mean aperiodic spectra for fully cleaned versus ocular contaminated data, with shaded areas indicating standard error of the mean. Panel E-J present results from the incremental contamination approach for blinks and saccades, using the same visualization schemes as B-D. Contamination levels reflect the proportion of artifactual segments mixed into clean data (clean: 0/15, low: 1/15, medium: 3/15, high: 5/15 segments), with the contamination effect topographies displaying standardized regression coefficients for the clean vs. medium contrast.

**Table 2:** Statistical results (CBPTs): Effect of physiological artifact contamination on the aperiodic offset and exponent.

| | Aperiodic offset<br>(avg. standardized $\beta$,<br>cluster p) | | Aperiodic exponent<br>(avg. standardized $\beta$,<br>cluster p) | |
| --- | --- | --- | --- | --- |
|  | Main dataset | Validation dataset | Main dataset | Validation dataset |
| **Ocular artifacts** |  |  |  |  |
| Component retention | $\bar{\beta} = 0.21$,<br>$p \leq 0.001$ | $\bar{\beta} = 0.28$,<br>$p \leq 0.001$ | $\bar{\beta} = 0.21$,<br>$p \leq 0.001$ | $\bar{\beta} = 0.22$,<br>$p \leq 0.001$ |
| Incremental contamination:<br>Blinks | $\bar{\beta} = 0.66$,<br>$p \leq 0.001$ | $\bar{\beta} = 0.88$,<br>$p \leq 0.001$ | $\bar{\beta} = 0.53$,<br>$p \leq 0.001$ | $\bar{\beta} = 0.74$,<br>$p \leq 0.001$ |
| Incremental contamination:<br>Saccades | $\bar{\beta} = 0.21$,<br>$p \leq 0.001$ | $\bar{\beta} = 0.36$,<br>$p \leq 0.001$ | $\bar{\beta} = 0.18$,<br>$p \leq 0.001$ | $\bar{\beta} = 0.28$,<br>$p \leq 0.001$ |
| **Muscular artifacts** |  |  |  |  |
| Component retention | $\bar{\beta} = -0.11$,<br>$p \leq 0.001$ | $\bar{\beta} = -0.09$,<br>$p \leq 0.001$ | $\bar{\beta} = -0.27$,<br>$p \leq 0.001$ | $\bar{\beta} = -0.24$,<br>$p \leq 0.001$ |
| Incremental contamination:<br>EEG based | $\bar{\beta} = -0.08$,<br>$p = 0.017$ | $\bar{\beta} = 0.05$,<br>$p = 0.086$ | $\bar{\beta} = -0.19$,<br>$p \leq 0.001$ | $\bar{\beta} = -0.17$,<br>$p \leq 0.001$ |
| Incremental contamination:<br>EMG based | $\bar{\beta} = 0.06$,<br>$p = 0.157$ | n.a. | $\bar{\beta} = -0.14$,<br>$p \leq 0.001$ | n.a. |
| **Cardiac artifacts** |  |  |  |  |
| Component retention | $\bar{\beta} = 0.03$,<br>$p \leq 0.001$ | $\bar{\beta} = 0.08$,<br>$p \leq 0.001$ | $\bar{\beta} = 0.02$,<br>$p = 0.04$ | $\bar{\beta} = 0.04$,<br>$p = 0.067$ |

*Note: For the incremental contamination approach, p values refer to the significance of the contamination factor in the linear mixed model. Standardized $\beta$ 's refer to the contrast "medium contamination vs. no contamination" and are averaged across all electrodes of the significant cluster. If no cluster was significant, $\beta$ 's are averaged across the largest non-significant cluster. n.a. indicates that the analysis was not applicable to the dataset.*

The incremental contamination approach further substantiated these findings. By leveraging co-registered eye-tracking data to identify ocular events, we separately assessed blinks and saccades, an analysis not possible with automated IC classification. Both blinks and saccades significantly increased the aperiodic offset and exponent, with blinks exerting a more pronounced influence (Table 2, Figure 3E-3J).

Finally, we tested the limits of standard denoising. Supplementary analyses (Supplement 5) demonstrated that blink and saccade-related distortions largely persisted even after removal of all ICs identified as artifacts. This suggests that standard preprocessing pipelines may not fully eliminate ocular influences, leaving them as potential confounds in neural interpretations of aperiodic parameters.

#### Muscular artifacts

In the component retention approach, muscular artifacts significantly influenced both the aperiodic offset and the aperiodic exponent across both datasets (see Table 2). Strikingly, in contrast to the findings for data quality and ocular artifacts, muscular contamination resulted in a significant decrease in the aperiodic offset and a flattening of the aperiodic component. These effects were not confined to specific electrodes but were observed broadly across the scalp (Figure 4B and 4D).

**Figure 4:**
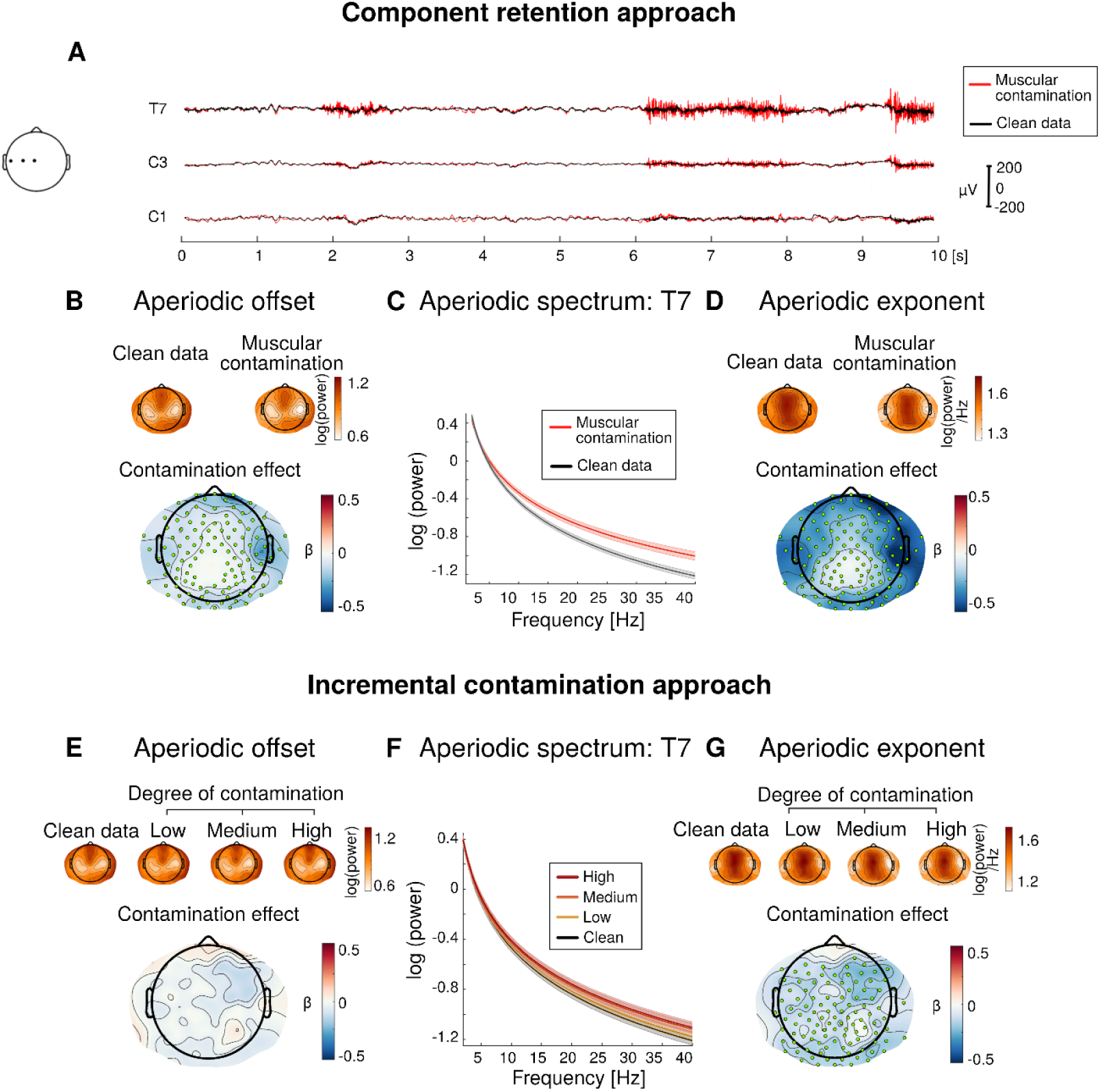
Impact of muscular artifact contamination on aperiodic parameter estimation. Visualizations represent the main dataset. Panels A-D illustrate findings from the component retention approach. Panel A shows prototypical data from three centrotemporal electrodes (indicated on head schematic from left to right: T7, C3 and C1) before (red) and after (black) the removal of ICs identified as muscular activity. Small topographies above B and D display aperiodic parameters derived from fully cleaned data (all artifactual ICs removed) and partially cleaned data (muscular ICs retained). Large maps show standardized regression coefficients from the linear mixed effects models predicting the aperiodic parameters from muscular contamination; green dots indicate electrodes significant in CBPT. Panel C shows mean aperiodic spectra for fully cleaned data versus data in which muscular ICs were retained, with shaded areas indicating the standard errors of the mean. Panels E-G presents results from the incremental contamination approach using direct EMG-artifact detection. Contamination levels reflect the proportion of artifactual segments mixed into clean data (clean: 0/15, low: 1/15, medium: 3/15, high: 5/15 segments), with the contamination effect topographies displaying standardized regression coefficients for the clean vs. medium contrast.

The incremental contamination approach substantiated these findings using two independent artifacts detection methods: An established EEG-based algorithm^34^ (both datasets) and EMG-based detection (main dataset only). We observed robust effects on the aperiodic exponent across detection methods (Table 2). Similar trends were observed for the offset (Figure 4E), though these reached statistical significance only with the EEG-based detection in the main dataset (Table 2; Supplement 6).

Consistent with ocular artifact results, the influence of muscular contamination on the aperiodic exponent persisted even when using the “fully cleaned” data, that is, when contaminated time windows were identified in basic preprocessed data, but aperiodic parameters were computed from the same time windows after ICA-based artifact removal (Supplement 5). This reinforced the concerns that well-established artifact correction methods may not be sufficient to prevent non-neural activity from masking neurobiological shifts in the aperiodic signal.

#### Cardiac artifacts

Findings regarding cardiac artifacts should be interpreted with caution due to two primary methodological constraints. First, the available sample sizes were smaller than in the other analysis (main dataset: N = 56, validation dataset: N = 35), as ICA did not identify cardiac ICs in a substantial proportion of participants (Supplement 1 & 2). Second, the incremental contamination approach could not be applied because typical resting heart rate of healthy adults (60 and 90 beats per minute^35^) precludes the extraction of artifact-free one-second segments.

In the component retention approach, cardiac artifact contamination was associated with a significant increase in the aperiodic offset across both datasets, with effects broadly distributed across the scalp (Figure 5B). Notably, this relationship was robust in within-participant analyses (Figure 5B) but absent in between-subject comparisons (Figure 5C). Regarding the aperiodic exponent, we observed a marginal positive relationship with cardiac contamination, although this did not reach the significance threshold of p<0.025 (Table 2).

**Figure 5:**
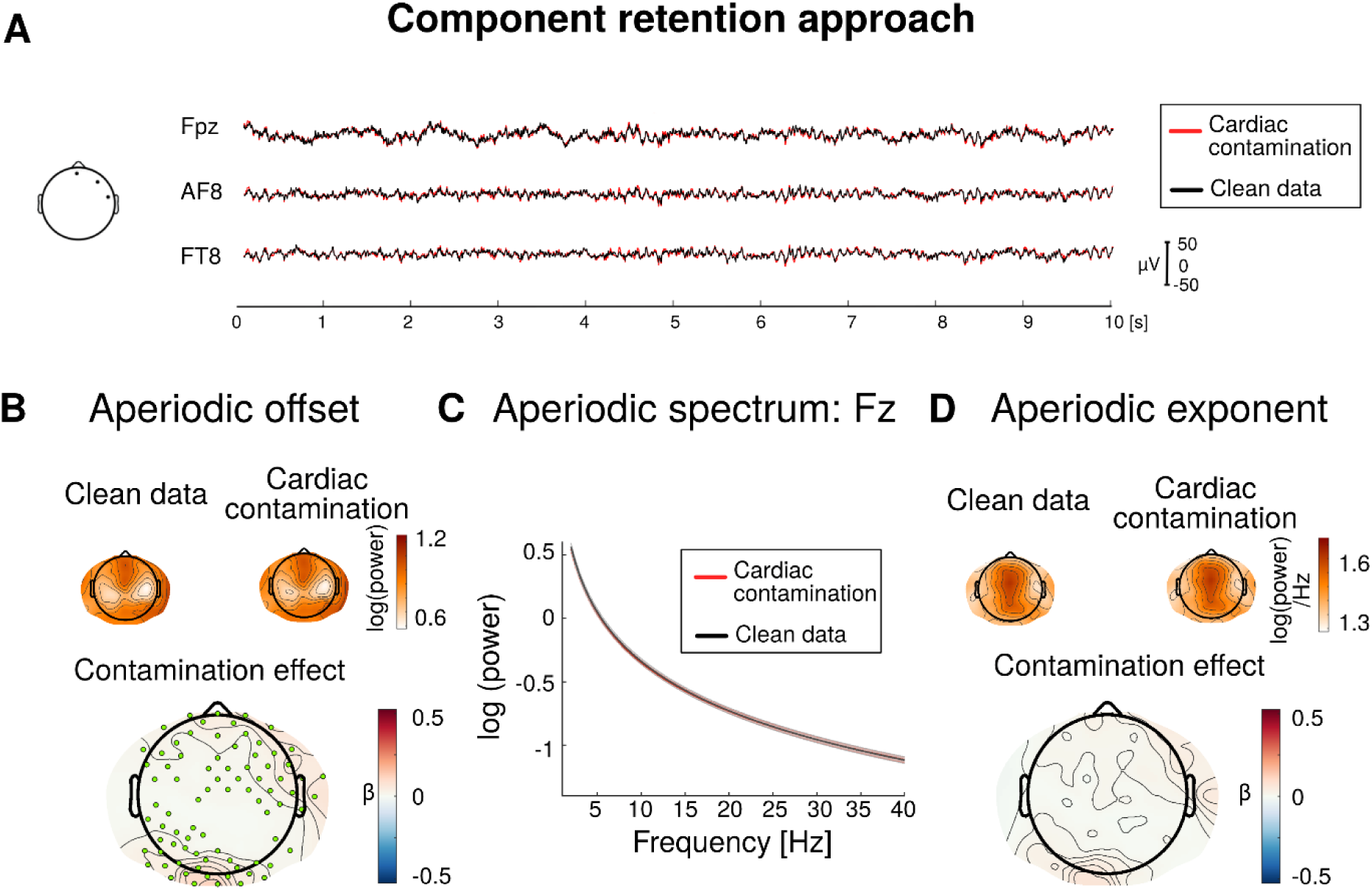
Impact of cardiac artifact contamination on aperiodic parameter estimation. Visualizations represent the main dataset. Panel A shows prototypical data from three frontotemporal electrodes (indicated on head schematic from midline to right: Fpz, AF8 and FT8) before and after removal of ICs identified as cardiac activity. Small topographies above B and D display aperiodic parameters derived from fully cleaned data (all artifactual ICs removed) and partially cleaned data (cardiac ICs retained). Larger maps below show standardized regression coefficients from the linear mixed effects models, predicting the aperiodic parameters from cardiac contamination; Green dots mark significant electrodes in CBPT. Panel C shows mean aperiodic spectra for fully cleaned data and of data in which cardiac ICs were retained, with shaded areas indicating standard errors of the mean.

### Result summary

Together, our systematic analyses demonstrate that both the aperiodic offset and exponent are highly sensitive to EEG recording quality and physiological artifacts. Figure 6 synthesizes these effects by presenting standardized β values averaged across all electrodes, providing an overall estimate of how each non-neural source impacts aperiodic estimation. The summary plot further highlights the directionality of the biases, methodological convergence across analytic approaches (component retention and incremental contamination), and consistency across independent datasets.

**Figure 6:**
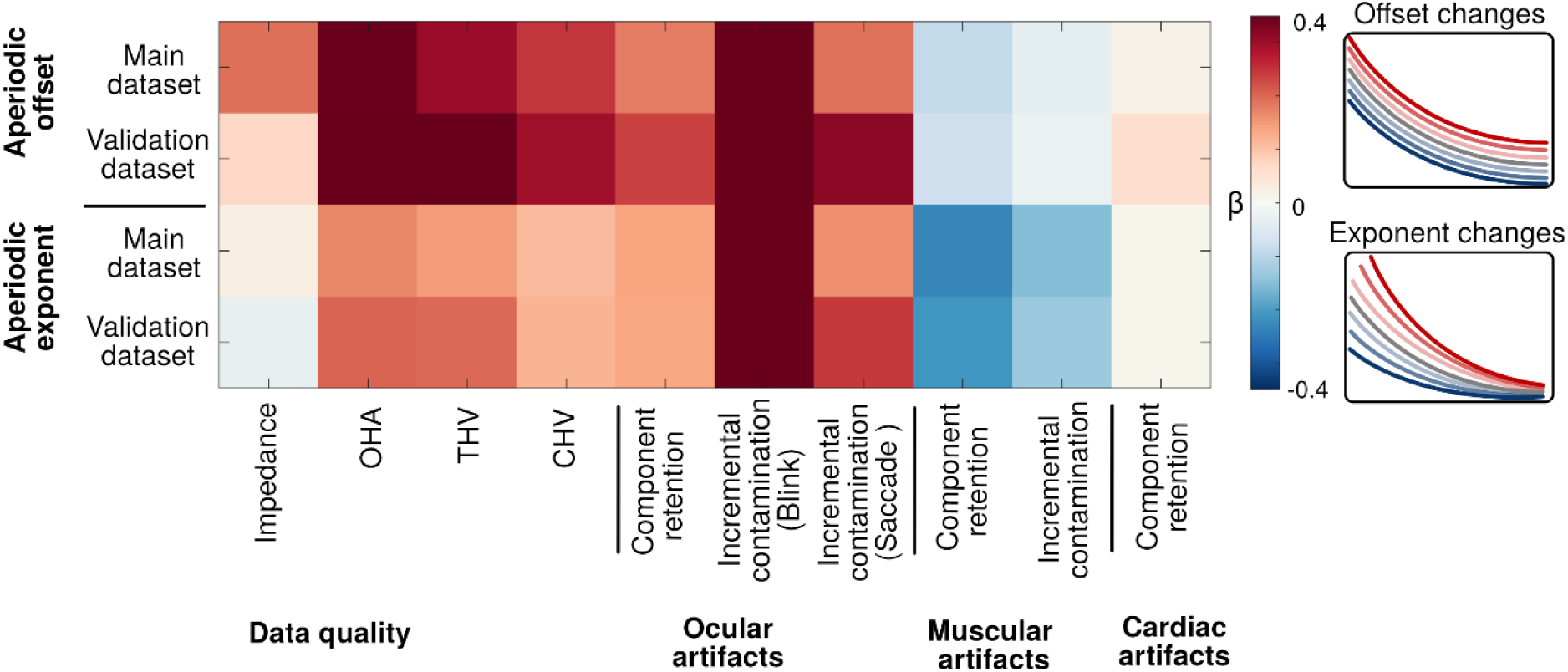
Synthesis of non-neural influences on aperiodic parameter estimation. Summary of standardized β coefficients averaged across all scalp electrodes for the main dataset and validation dataset. For the component retention approach, β estimates represent the contrast between data with artifactual ICs retained versus removed. For the incremental contamination approach, β estimates reflect the contrast between medium contamination (3 out of 15 segments) and clean data. Note that average β estimates for incremental blink contamination reached magnitudes up to 0.87, while OHA and THV metrics reached up to 0.50, highlighting the substantial impact of these sources on spectral estimates. Schematic plots (right) illustrate how these reported beta estimates correspond to changes in each aperiodic parameter.

### Mitigating artifact-induced biases through statistical correction

To provide a practical solution for scenarios where populations or experimental conditions differ systematically in artifact rates (e.g. comparing typical developing children and children with an ADHD diagnosis^31^), we conducted an additional simulation analysis using the main dataset. We modeled a plausible scenario comparing populations with different levels of ocular contamination: 20% artifactual segments, representative of healthy populations^27^, versus 40%, typical in clinical samples^27,31^. For illustrative purposes, we focused on a single artifact type (blinks) and electrode (POz), and on the aperiodic exponent, as it is more commonly studied than the offset^9,17^.

The main dataset was randomly divided into two subgroups, and differential eye-blink contamination was introduced using the incremental contamination approach (Figure 7A). Subgroup 1 consistently received low-contamination load (mean 3 contaminated segments of 15 total). Subgroup 2 was sampled twice, once under low-contamination condition (Group 2A, identical to Subgroup 1) and once under high-contamination condition (Group 2B, mean 6 of 15 segments). Aperiodic exponents were extracted at electrode POz from “fully cleaned” data (i.e., after removal of all artifactual ICs).

**Figure 7:**
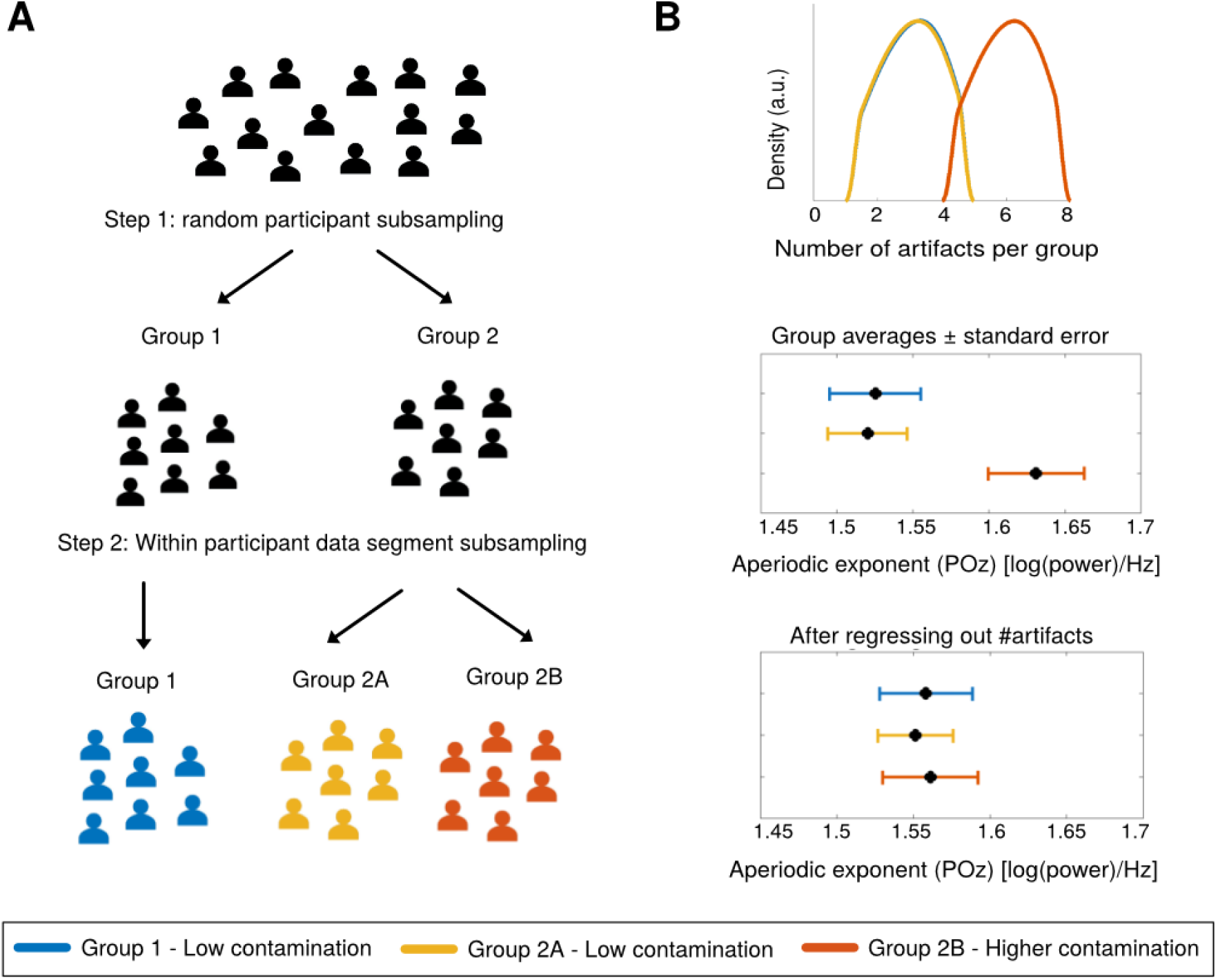
Statistical mitigation of artifact-induced biases in aperiodic parameter comparisons. Example shown for the effect of eye blinks on the parieto-occipital aperiodic exponent, based on “fully cleaned” data (all artifact ICs removed). Panel A shows the random subsampling of participants in the main dataset. Panel B displays artifact segments per subgroup (top) and the resulting aperiodic exponent estimates (bottom) plotted as group means ± standard error, before (upper) and after (lower) regressing out the number of artifacts.

As expected, groups with matched contamination levels (Group 1 vs. Group 2A) showed no significant difference in aperiodic exponents (standardized β = 0.05, p = 0.82). However, the comparison between low- versus high-contamination groups yielded a significant, spurious group difference (Group 1 vs. Group 2B, standardized β = 0.54, p = 0.01, Figure 7B, middle panel). Crucially, after regressing out the number of contaminated segments (residualizing the aperiodic exponent by contamination level), this spurious group difference was eliminated (Group 1 vs Group 2B, standardized β = 0.08, p = 0.69, Figure 7B, bottom). A permutation-based robustness analysis (5000 iterations) confirmed that this pattern was stable and not dependent on a specific random split (see Supplement 12 for a distribution of resulting p and standardized β values).

These results demonstrate that regression-based statistical correction effectively mitigates artifact-induced biases in aperiodic parameter comparisons. When populations or conditions differ systematically in artifact rates, quantifying and statistically controlling for these differences enables valid neurobiological inference while maintaining statistical power.

## Discussion

Our comprehensive analyses revealed that both data quality and physiological artifacts substantially affect aperiodic parameter estimation. Distinct directional patterns emerged across artifact types: poor data quality, ocular activity, and cardiac artifacts were associated with larger offsets and exponents (i.e., steeper aperiodic slopes), whereas muscular artifacts produced the opposite pattern (i.e., smaller offsets and flatter aperiodic slopes). Importantly, these distortions were not confined to local regions but extended across the entire scalp, indicating that artifact sources influence aperiodic estimates even at distant electrodes. Crucially, these effects persisted after applying state-of-the-art preprocessing at realistic contamination levels, suggesting that typical artifact correction procedures may not fully remove non-neural contributions from aperiodic parameters. Given that the magnitude of these influences is comparable to group differences often interpreted as physiologically meaningful, our findings underscore a significant risk of spurious neural inference in current literature.

The mechanisms underlying these effects likely stem from how specific spectral signatures interact with parameter estimation algorithms. As pointed out by a previous study^36^, oscillatory peaks near the lower or upper bound of the fitted frequency ranges can bias the estimation of the aperiodic parameters in the algorithm of specparam^1^, but also in an alternative estimation method such as irregular-resampling autospectral analysis (IRASA^37^). Because non-neural artifacts typically exhibit high amplitudes, they are particularly susceptible to such misestimation.

Our findings validate this framework: Ocular artifacts introduce high-amplitude low-frequency noise into the EEG signal^38,39^, primarily affecting the lower end of the frequency spectrum used for aperiodic fitting (2-40 Hz) and thereby increasing both the aperiodic offset and exponents. Conversely, muscular artifacts introduce high-frequency noise^34^, elevating power near the upper bound of the fitting range (∼40Hz) and consequently resulting in smaller offsets and exponents. Poor data recording quality often reflect elevated low-frequency noise^40^, yielding larger aperiodic offsets and steeper aperiodic signal components. Together, these results highlight that aperiodic parameter estimation is vulnerable to systematic bias when non-neural sources introduce spectral power that interferes with typical fitting ranges applied to the algorithm, challenging the presumed independence of periodic and aperiodic components.

Therefore, our findings have profound implications for basic and clinical research. While the component retention approach confirms that ICA substantially reduces contamination, significant residual influences persist. This poses a major risk for comparative research: observed aperiodic differences may mistakenly be attributed to neural mechanisms when they in fact reflect systematic group variations in artifact prevalence. This concern is supported by prior work showing that observed age-related differences in aperiodic parameters were linked to cardiac activity, and that ICA-based rejection did not fully remove these effects^29^. Our findings extend this observation, demonstrating that data quality and all major physiological artifact types (ocular, muscular, and cardiac) challenge the validity of interpreting uncorrected group differences as shifts in the excitation-inhibition (E/I) balance.

It is therefore imperative to determine whether study groups or experimental conditions differ systematically in data quality and artifact contamination. Across neuroimaging modalities, artifacts are rarely distributed equally across populations. For example, greater in-scanner head motion has been reported in ADHD^30,41^, in ASD^30^ and in schizophrenia^30^ populations. Also EEG studies have found higher artifact rates in children and adolescents compared to adults, with further increases in pediatric ADHD^31^, and elevated ocular artifacts have been demonstrated in schizophrenia patients compared to controls^42^. However, as demonstrated in our simulation (Figure 7), statistical control of contamination levels effectively eliminate these biases. By regressing out artifact prevalence, researchers can disentangle non-neural noise from genuine neurobiological signals, ensuring more robust and reproducible between-group comparisons.

While our findings provide systematic insights into the influence of artifacts on aperiodic parameter estimation and how such effects can be mitigated, several methodological limitations should be acknowledged. First, the efficacy of statistical correction depends on accurate artifact identification. While we leveraged eye-tracking and EMG ground truth, many extant datasets lack these auxiliary measures. Developing standardized artifact detection protocols therefore represents a crucial step for the field.

Second, although ICA-based artifact correction is standard practice, components labeled as artifacts may also contain neural signals, potentially biasing results from the component retention approach. This risk may be amplified when using automated classifiers (e.g., ICLabel), even though automation enhances objectivity and reproducibility. However, the strong convergence with our incremental contamination analyses, which avoided ICA-based removal entirely, suggests that any potential neural signal loss did not drive the observed effects.

Finally, our analyses focused on the specparam algorithm, which may limit generalizability to alternative aperiodic parameterization methods. However, specparam was chosen because it is the most widely used approach for aperiodic parameter estimation in the field^9,17^. Other approaches, such as IRASA, rely on different modeling principles and may show some method-specific variation.

Looking ahead, future studies should extend our findings to alternative parametrization approaches and beyond resting-state data, assessing artifact influences across diverse experimental paradigms, e.g. when tracking temporal dynamics of the aperiodic signal as proposed in Wilson et al^43^. Most importantly, the field must systematically re-evaluate previously reported aperiodic findings to determine whether observed developmental trajectories or clinical group differences remain significant after controlling for differential artifact rates.

In conclusion, our findings demonstrate that data quality and physiological artifacts introduce systematic biases in aperiodic signal parameters, with magnitudes comparable to those often interpreted as meaningful neurophysiological effects. While state-of-the-art preprocessing can attenuate these biases, they remain sufficiently pronounced to influence reported group or condition differences. As aperiodic parameters gain prominence in studies of brain function across development, aging, and clinical populations, recognizing and statistically controlling for these contamination sources will be essential for the valid characterization of the human brain’s aperiodic signal component.

## Methods

### Datasets

All analyses were conducted on a main and an independent validation dataset to enhance the robustness of our results, and to evaluate their generalizability across different EEG systems. In both datasets, EEG and eye tracking data were collected simultaneously during resting state recordings. In the main dataset, which was specifically recorded for the purpose of this study, additional ECG and EMG channels were included to improve the identification of physiological artifacts. The following sections describe each dataset and the corresponding EEG hardware used for data acquisition.

#### Main dataset

The main dataset comprised 99 healthy young participants aged 18-35 years. Individuals were excluded if they exhibited psychiatric symptoms, suffered from severe neurological disorders or if they currently used psychotropic drugs such as antidepressants, alpha-agonists, neuroleptics and mood stabilizers, or recreational drugs. The study was approved by the ethics committee of the Faculty of Arts and Social Sciences of the University of Zurich, and all participants provided written informed consent.

Main dataset: Data acquisition

For the main dataset, EEG was acquired using a 128-channel ANT Neuro system (ANT Neuro, Hengelo, Netherlands) with a sampling rate of 500Hz. Recordings were referenced to CPz, and impedances were maintained below 20 kΩ. In addition to the EEG channels, a horizontal and vertical electrooculogram (EOG), three bipolar EMG channels, and a two-electrode ECG were obtained with the same sampling rate of 500Hz. The EMG electrode pairs were placed above the left eyebrow, on the left cheek (above the masseter muscle), and on the right upper neck (to capture activity of the trapezius and the sternocleidomastoid muscle). Electrode locations were selected following prior work capturing EMG activity in the context of EEG recordings^44^. For the ECG, one electrode was placed in the right infraclavicular area (below the clavicle at the midclavicular line) and the other on the left lower torso, near the iliac crest. EEG and ECG were recorded with a common ground, positioned on the right mastoid. Eye movements were simultaneously recorded by an infrared video-based eye tracker (EyeLink 1000 Plus, SR Research) at a sampling rate of 500 Hz and an instrumental spatial resolution of 0.01°.

#### Validation dataset

The second employed validation dataset was previously recorded in our laboratory and comprised 103 healthy participants aged 18–35 years. The same exclusion criteria as for the main dataset were applied here. The study was approved by the Institutional Review Board of Canton Zurich (BASEC-Nr. 2017–00226), and all participants provided written informed consent.

Validation dataset: Data acquisition

EEG data was recorded at a sampling rate of 500 Hz using a 128-channel EEG Geodesic Sensor Net system (Electrical Geodesics, Eugene, Oregon). Recordings were referenced to Cz (vertex of the head), and electrode impedances were kept below 40 kΩ. Additionally, eye movements were recorded by the same infrared video-based eye tracker as in the main dataset (EyeLink 1000 Plus, SR Research).

### Experimental setup and procedure

Identical procedures were used for the collection of both datasets. Participants were seated in a sound- and electrically shielded Faraday cage, equipped with a 24-inch screen (ASUS ROG Swift PG248Q, resolution 800 × 600 pixels) and a chinrest positioned 68cm from the screen. Resting-state recordings were obtained immediately after electrode preparation (i.e., impedances were below 20 kΩ in the main dataset or 40 kΩ in the validation dataset) and the calibration of the eye tracker. The eye tracker calibration and validation were performed on a 9-point grid until the average validation error across all points was <1°.

Resting-state EEG data was recorded while participants alternated between eyes-open and eyes-closed conditions. Audio cues delivered through in-cage loudspeakers indicated the transitions between the conditions. Following an established protocol^45,46^, participants completed five cycles, each consisting of 20 s of eyes-open fixation followed by 40 s of eyes-closed rest, yielding a total of 100 s of eyes-open and 200 s of eyes-closed data for analysis.

### Preprocessing

The same preprocessing pipeline was applied to the full resting-state recordings of both datasets, including the alternating eyes-open and eyes-closed periods. In the main dataset, the auxiliary electrodes (EOG, ECG, EMG) were excluded from the below described preprocessing steps to preserve their raw signals, however they were included for the ICA decomposition to improve the identification of ocular, cardiac and muscular artifact ICs.

Because the primary focus of this study was to examine the influence of data quality and physiological artifacts on estimates of the aperiodic signal, preprocessing followed a state-of-the-art pipeline to mirror realistic research practice. Therefore, the automated preprocessing toolbox Automagic^47^ was employed, which performed detection and interpolation of bad channels, a 0.5 Hz high pass filter and line noise removal (“Basic preprocessing“; Figure 1).

Independent component analysis (ICA) was then performed. To allow for subsequent analyses of selective artifact removal and retention (see below), all ICs were retained in the data. For the pipeline’s automated data-quality assessment, however, artifactual ICs (classified by ICLabel) were temporarily removed. Datasets rated as poor quality were excluded from further processing (see Supplement 7 for a detailed description of the pipeline). Before the final extraction of the aperiodic signal component, data was rereferenced to the common average reference.

For all analyses, except the incremental contamination analysis on ocular artifacts (see below), data from the eyes-closed condition were used, which is most commonly employed in resting state EEG studies^48^. Therefore, data of the five 40-second segments were concatenated. The first and last two seconds of each segment were excluded to remove potential artifacts of the acoustic prompts and of eyes closure or eyes opening, respectively, yielding a total of 180 seconds of eyes-closed data. For the eyes-open data, a total of 80 seconds was available after merging and applying the same exclusions.

### Calculation of the aperiodic signal component

For all analysis approaches outlined below, the aperiodic parameters were calculated as follows. The power spectral density (PSD) was first computed using Welch’s Method^49^, with a window length of 1 second. The resulting PSD was then parameterized using the specparam algorithm^1^, which decomposes the power spectrum into an aperiodic component and oscillatory peaks. In brief, the algorithm iteratively models the PSD as an aperiodic component (L) and a set of Gaussians representing oscillatory peaks rising above the aperiodic signal. A detailed description of the fitting procedure is provided in Donoghue et al.^1^

The aperiodic component is defined as:

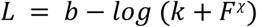

Here, b represents the aperiodic offset, F, the vector of input frequencies, and χ the aperiodic exponent. The additional k parameter is set to 0 in the present analysis, as this parameter allows to model a bend of the aperiodic component which is primarily needed when fitting signals in broad frequency ranges, typically in intracranial recordings^2^. Further settings for the algorithm were: peak width limits: [1, 8]; max number of peaks: infinite; minimum peak height: 0; peak threshold: 2 sd above mean. The model was fitted to a frequency range of 2-40 Hz, consistent with the original specparam algorithm publication^1^. Furthermore, this range was commonly used in subsequent studies examining the relationship between aperiodic activity and age^12–14,50,51^ and was most frequently applied in clinical investigations on aperiodic parameters^9^.

Across all analytic subsamples, the specparam algorithm achieved consistently high model fits across (R² = 0.96-0.99), regardless of differences in EEG data quality and artifact contamination levels between approaches (see Supplement 8 for a detailed assessment of model fits across analyses).

### Effect of data quality on the aperiodic signal

As depicted in Figure 1 (green box), analyses of the relationship between data quality and the aperiodic signal parameters were conducted on fully cleaned data (i.e. after removal of all artifactual ICs). This approach was chosen to mirror a realistic scenario, enabling us to assess whether the quality of data acquisition influences the estimated aperiodic parameters even after state-of-the-art preprocessing.

#### Hardware based quality metrics: Electrode impedances

As a first measure of recording quality, electrode impedances were extracted. In both study protocols (i.e. in the main and the validation dataset), impedances were recorded immediately after electrode preparation by the research assistants, and resting-state data collection began directly thereafter. Therefore, impedance measures corresponded to the beginning of the analyzed EEG recordings. For subsequent statistical analyses, only impedance values within ranges commonly accepted in EEG research were included. Following the recommendation of the EEG system manufacturers, the cut-off was set to 20 kΩ for main dataset (electrolyte gel-based EEG system), and to 50 kΩ for the validation dataset (salt water-based EEG system). The distribution of electrode impedances is shown in Supplement 9.

#### Algorithmic quality metrics

The second analysis examined the relationship between the aperiodic signal parameters and objective, algorithmic estimates of data quality, assessed at the end of the preprocessing pipeline (i.e.after temporal removal of all artifactual ICs). Whereas the thresholded version of these quality metrics were used during preprocessing to exclude bad quality data (see section “Preprocessing” and Supplement 7), we here used their continuous values. In more detail, these metrics were defined as described and validated in Pedroni et al.^47^: Ratio of data with overall high amplitude (OHA), defined as the ratio of datapoints (i.e. channels × timepoints) that have an absolute voltage value larger than 20 μV. Ratio of timepoints of high variance (THV) measures the ratio of time points where the standard deviation across channels exceeds 10 μV. Ratio of channels of high variance (CHV), the final measure, is defined as the ratio of channels whose standard deviation across all timepoints exceeds 10 μV. Unlike electrode impedances, which are measured individually for each electrode, these three algorithmic quality metrics each yield a single summary value for the entire EEG recording. Distributions of the three quality metrics are provided in Supplement 9.

Relationships among all quality measures are evaluated in Supplement 10. In brief, electrode impedances and algorithmic metrics were largely uncorrelated and thus reflect distinct aspects of data quality, whereas the three algorithmic metrics showed high intercorrelations, indicating substantial overlap, but still accounting for partly distinct variance components.

### Effect of physiological artifacts on the aperiodic signal component

The second set of analyses investigated the influence of physiological artifacts on the estimated aperiodic parameters (Figure 1, blue boxes). Two complementary approaches were employed: *component retention* and *incremental contamination*. The following sections describe each in more detail.

#### Component retention approach

This approach assessed the impact of physiological artifact contamination by comparing the aperiodic signal estimated on EEG data while either retaining or removing ICs representing non-neural activity. Probability scores assigned by ICLabel during preprocessing were used for classification. In a first step, all ICs reflecting artifacts (ocular, muscular, cardiac, line noise, channel noise) with probabilities >0.8 were removed from the data and the aperiodic signal was estimated based on this cleaned dataset (i.e. “fully cleaned data”). Next, to isolate the effect of specific artifact types, the IC removal procedure was repeated in three separate iterations. In each iteration all artifactual ICs were removed except those of a single target class (e.g., ocular ICs retained; see Fig. 1 for a visualization). After each selective retention, the aperiodic signal was re-estimated, thereby enabling assessment of the influence of that artifact category on the aperiodic parameters. Supplement 2 visualizes the number of ICs for each artifact category per participant.

In line with typical resting-state paradigms, these analyses were conducted on the merged eyes-closed EEG data. Importantly, eye movements continue to occur under lid closure, even when participants are instructed to maintain a given position of the eye^52^. However, for supplementary analysis, the effect of ocular artifact contamination was also estimated on eyes-open resting state data.

#### Incremental contamination approach

Because ICA cannot perfectly separate neural from non-neural signals, we implemented an alternative approach to complement the component retention approach. This second approach was designed to reduce the risk that removing ICs identified as artifacts would also eliminate neural activity and thereby affect the estimated aperiodic parameters. Consequently, instead of relying on ICA-based artifact correction, the aperiodic parameters were estimated either from entirely artifact-free segments only, or from segments that intentionally included varying levels of artifact contamination. In a first step, two algorithms (described below) were applied to the basic preprocessed resting-state EEG data (i.e. prior to IC removal) to identify segments containing physiological artifacts. In a next step, 15 artifact-free one-second segments were extracted, and the aperiodic parameters were calculated on these clean segments. Subsequently, either one, three or five of the clean segments were exchanged with artifact-contaminated segments of the same length, and the aperiodic parameters were recalculated for each configuration (see Figure 1 for a visualization). Only participants were included who had at least 15 clean segments, and five segments containing artifacts respectively. The number of 15 clean segments was chosen to balance participant retention with sufficient model fit quality for the specparam algorithm (see Supplement 2). The chosen contamination levels (∼7%, 20%, and 33%) reflect typical ranges of artifact contamination in EEG data, which often vary between 20–40% depending on the investigated population^27,31^. The full distribution of the number of available segments per subject is presented in Supplement 2.

For supplementary analyses, artifact-contaminated time windows (identified in the basic preprocessed data) were used to extract corresponding segments from the fully cleaned EEG (i.e., after removal of all artifactual ICs, as described above). This allowed us to assess whether residual effects of physiological artifacts on the aperiodic parameter estimation persisted after state-of-the-art artifact correction.

The following two sections describe how segments containing ocular and muscular artifacts were determined.

#### Ocular artifacts

To examine the influence of saccades and blinks on the aperiodic signal parameters, the incremental contamination approach was applied to merged eyes-open resting-state data. Using eyes-open data allowed us to incorporate eye tracking information as a ground truth to identify segments containing ocular artifacts. EEG and eye-tracking data were co-registered with the EYE-EEG toolbox^53^. Artifact-free segments were defined as one-second EEG epochs during fixation periods (i.e., during periods of no eye movement) as indicated by the eye tracker. From these, 15 segments were randomly selected. Segments containing blinks were identified by epoching around blink onsets (–200 to +800 ms), detected by the eye tracker. From all potential segments containing a blink, either one, three or five segments were randomly selected. Likewise, segments containing saccades were determined by segmenting around saccade events.

#### Muscular artifacts

Muscular artifacts during merged eyes-closed resting-state recordings were identified using two complementary algorithms. The first artifact detection algorithm was based on EEG data and thus could be applied to both the main and the validation dataset. To obtain a more direct measure of muscular activity, three bipolar EMG channels were additionally recorded in the main dataset (see above). Based on these electrodes, a second algorithm was implemented to identify segments containing muscular artifacts.

The EEG-based muscular artifact detection was performed using a state-of-the-art procedure^34^. Specifically, the first principal component of the EEG data, which typically contains high amplitude artifacts^54^, was extracted. This component was then filtered to higher frequency bands (65-115Hz) to enhance muscular contributions, and its envelope was low pass filtered (4Hz) to remove transient noise peaks. This signal was finally thresholded to detect artifact segments (see Supplement 11 for a detailed description).

For the main dataset, EMG-based muscular artifact detection was performed on the three bipolar EMG channels. Signal envelopes were computed for each EMG channel (also low-passed filtered at 4 Hz to suppress transient noise peaks) and thresholded to detect muscular activity whenever any of the three processed channels exceeded the criterion (see Supplement 11 for details).

### Statistical analysis

Prior to statistical analyses, all continuous variables were z-transformed to standardize the regression coefficients, enabling direct interpretation of the resulting estimates in terms of effect size. To account for the problem of multiple comparisons across electrodes (105 or 128 channels, depending on dataset), cluster-based permutation tests (CBPTs) as implemented in FieldTrip^55^, in combination with custom-written functions, were employed.

#### Data quality: Linear models & cluster based permutation tests

To statistically evaluate the relationship between the aperiodic l parameters and the measures of data quality (i.e. electrode impedances and the algorithmic quality metrics CHV, THV and OHA), we fitted the following linear models:

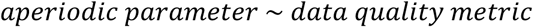

Note that the models were fitted for each quality metric independently and separately for the aperiodic offset and exponent. Spatial clusters were defined by thresholding the t-values of the β coefficient at a two-tailed p < 0.05 (i.e., t-values exceeding the critical t for p = 0.025 in each tail), and grouping neighboring electrodes showing significant t-values. Next, the t-statistics within each cluster were summed to calculate cluster-mass statistics (“maxsum” method). To generate the null distribution, 1000 permutations were performed by randomly shuffling data quality values across participants; for each permutation, the maximum cluster mass of the largest identified cluster was retained. A cluster in the observed data was deemed significant if its mass exceeded the 97.5th percentile of the permutation distribution of cluster masses (cluster-level α = .025, adjusted for two tails).

#### Physiological artifacts: Linear mixed effects models & cluster based permutation tests

To evaluate whether physiological artifact contamination influenced the estimated aperiodic signal parameters, similar CBPTs statistics were applied using linear mixed-effects models. Note that separate models were fitted for the aperiodic offset and the aperiodic exponent. For the component retention approach, the following model was applied:

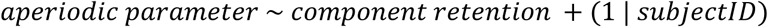

where the factor “component retention” indicated whether artifact-related ICs were removed (0) or retained (1). In this case, statistical inference was based on the fixed-effect t-statistics associated with the “component retention” predictor as provided by the linear mixed-effects model. These t-statistics were used as the input for cluster formation in the CBPT.

For the incremental contamination approach, the model was defined as:

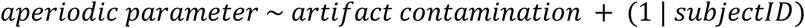

Here, the “artifact contamination” factor had four levels: 0 (clean data segments only), 1 = low contamination (1 artifactual segment), 2 = medium contamination (3 artifactual segments) and 3 = high contamination (5 artifactual segments). Random intercepts for subjects were included in both models to account for within-subject dependencies across repeated conditions.

For the incremental contamination analysis, test statistics were derived using a model comparison framework. Specifically likelihood ratio tests were conducted between each full model and a reduced baseline model:

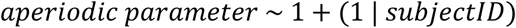

This model comparison approach was used to obtain a single test statistic reflecting the overall effect of the artifact factor, which can be employed for cluster formation, rather than performing separate CBPTs for each individual factor level. The likelihood ratio test statistic approximates a chi-square distribution, therefore, cluster formation is based on extreme values in this distribution at an alpha level of 0.05. Note that the likelihood-ratio test evaluates only whether the full model with the additional factor (i.e “artifact contamination”) provides a significantly better fit than the reduced model. Thus, this test evaluates a single directional hypothesis (improvement in fit), and therefore is a one-sided test (i.e. the α level of 0.05 was not further divided).

As described for the analysis on data quality, the sum of the test statistics in each formed cluster is saved for the observed data, and compared to a null distribution based on 1000 permutations of randomly shuffled factor levels within participants. Again, a cluster was deemed significant if its observed mass exceeded the 97.5th percentile of the permutation distribution (cluster-level α = .025, adjusted for two tails).

## Data availability

All data that support the findings of this study are available on: https://gin.g-node.org/mtroen/NonNeuralAperiodic

## Code availability

All analysis code is available on a public repository: https://gin.g-node.org/mtroen/NonNeuralAperiodic

## Author contributions

M.T. contributed: Writing – original draft, Methodology, Formal analysis, Data curation, Conceptualization

N.L. contributed: Writing – review & editing, Supervision, Project administration, Methodology, Formal analysis, Conceptualization.

## Acknowledgement

This work was supported by the Swiss National Science Foundation [100014_175875].

