## Supplementary materials for "Non-Neural Sources Systematically Impact Aperiodic EEG Activity"

### Supplement:

S1. Sample description including analytic subsamples
S2. Number of independent components and data segments of each artifact category per participant
S3. Side by side visualizations
S4. Component retention approach: Ocular artifact removal in eyes-open data
S5. Incremental contamination approach applied to fully cleaned data
S6. Incremental contamination approach: EEG-based muscle detection in main dataset
S7. Detailed preprocessing pipeline
S8. Specparam model fits per analysis
S9. Distribution of impedances and algorithmic quality metrics
S10. Relation between algorithmic quality metrics and electrode impedances
S11. Muscular artifact detection algorithm
S12. Robustness analysis of the statistical correction approach

##### S1. Sample description including analytic subsamples

Supplementary Table 1 presents sample sizes for each analysis. Samples varied depending on electrode impedance availability and successful artifact identification. The main dataset had more complete data coverage than the validation dataset.

**Supplementary Table 1**

*Participant demographic characteristics of the full sample and analytic subsamples*

| | Main dataset | | | Validation dataset | | |
|---|---|---|---|---|---|---|
| | N | mean age (sd) | sex (m/f) | N | mean age (sd) | sex (m/f) |
| Total | 89 | 22.9 (3.8) | 22/67 | 93 | 23.8 (3.0) | 39/54 |
| **Per analysis** | | | | | | |
| Electrode impedance | 89 | 22.9 (3.8) | 22/67 | 55 | 23.2 (2.3) | 28/27 |
| Algorithmic quality metrics | 89 | 22.9 (3.8) | 22/67 | 93 | 23.8 (3.0) | 36/57 |
| Component retention: Ocular artifacts | 88 | 23.0 (3.8) | 22/66 | 93 | 23.8 (3.0) | 36/57 |
| Component retention: Muscular artifacts | 86 | 22.9 (3.8) | 21/65 | 93 | 23.8 (3.0) | 36/57 |
| Component retention: Cardiac artifacts | 56 | 23.0 (3.6) | 14/42 | 35 | 23.6 (2.8) | 21/14 |
| Incremental contamination: Blink artifacts | 76 | 22.8 (3.8) | 17/59 | 86 | 23.9 (3.0) | 33/53 |
| Incremental contamination: Saccade artifacts | 55 | 23.3 (4.1) | 18/37 | 57 | 24.3 (3.1) | 22/35 |
| Incremental contamination: Muscular artifacts (EEG) | 63 | 22.2 (3.5) | 13/50 | 69 | 23.9 (3.1) | 25/44 |
| Incremental contamination: Muscular artifacts (EMG) | 75 | 22.9 (3.8) | 18/57 | n.a. | n.a. | n.a. |

*Note: Muscular artifact detection based on EMG data was not applicable (n.a.) to the validation dataset as no EMG channels were collected during its data acquisition.*

**S2. Number of independent components and data segments of each artifact category per participant**

Supplementary Figure 1 visualizes the number of independent components available per participant for the component retention approach. Participants with zero components of a target artifact category were excluded from the corresponding analysis.

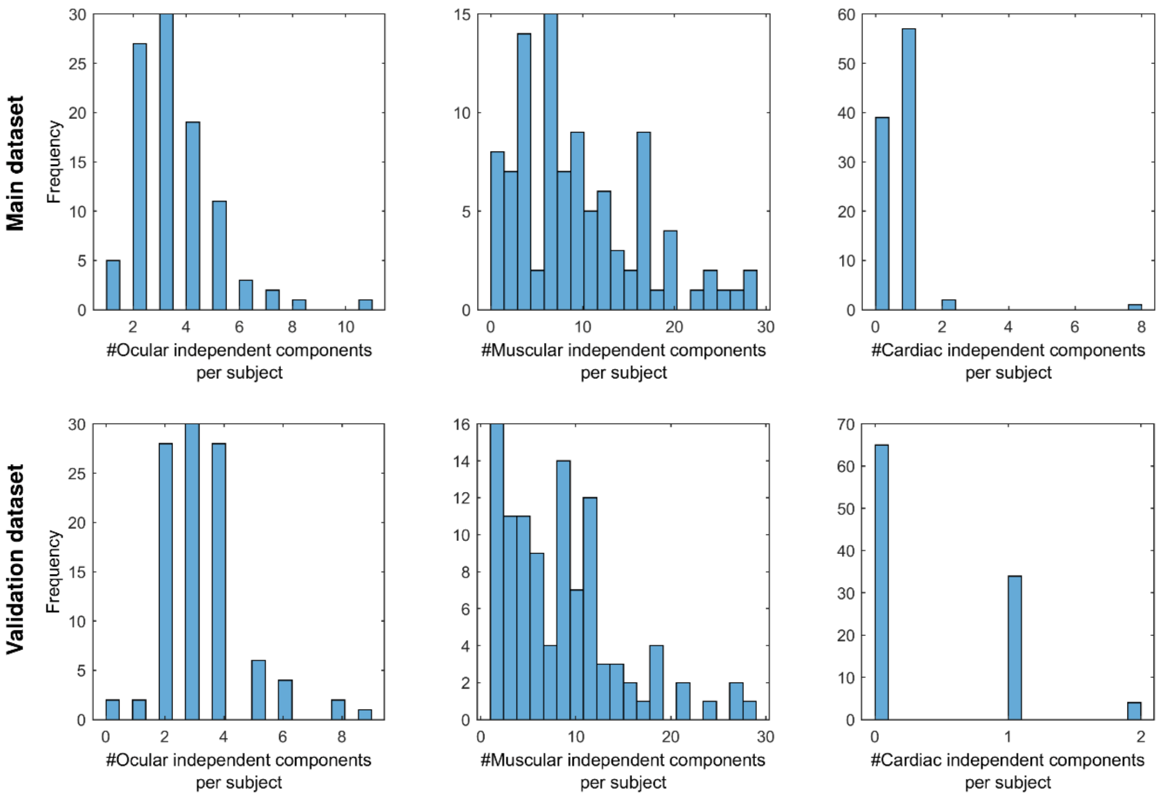


*Supplementary Figure 1: Number of independent components reflecting the three different artifact categories (ocular, muscular, cardiac) as labeled by ICLabel*[1].

Supplementary Figure 2 visualizes the number of available clean and contaminated data segments per participant for the incremental contamination approach.

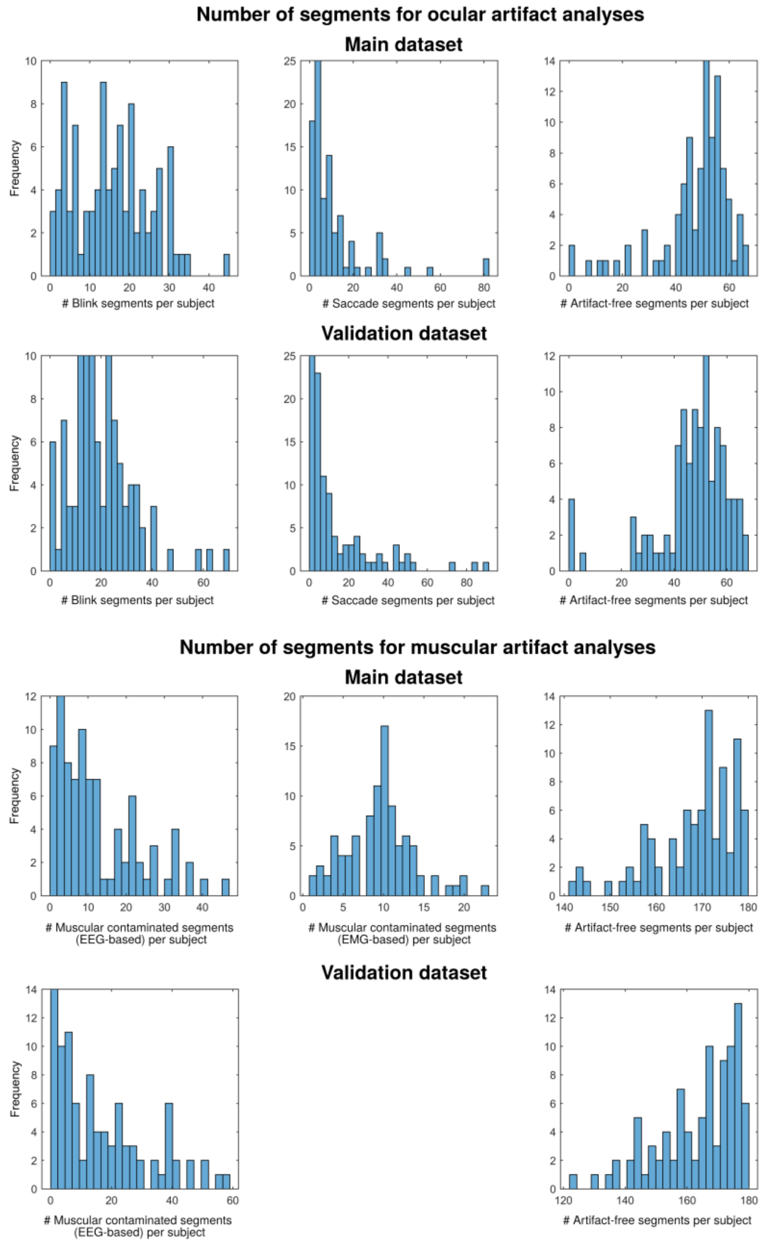


*Supplementary Figure 2: Rows A and C: Number of detected ocular artifacts (annotated by the eye tracker) and clean one-second segments (no annotated ocular activity) available for the incremental contamination approach investigating ocular contamination (eyes-open data). Note that one-second segments were epoched around each detected ocular artifact and could thus overlap in time. Rows B and D: Number of muscular-contaminated and clean one-second data segments (no muscular activity, eyes-closed data). For each analysis, 15 clean and 1, 3 or 5 of the contaminated segments were randomly selected. For this visualization purposes, two participants exerting more than 100 saccades, as well as two outliers showing more than 100 blinks were removed.*

#### S3. Side by side visualizations

Supplementary Figures 3-6 visualize all main results (i.e. impact of data quality, ocular, muscular and cardiac artifacts on the aperiodic parameters) of the main and the validation dataset for direct visual comparison. Note that statistical results are described in detail in the main manuscript (Table 1 and 2).

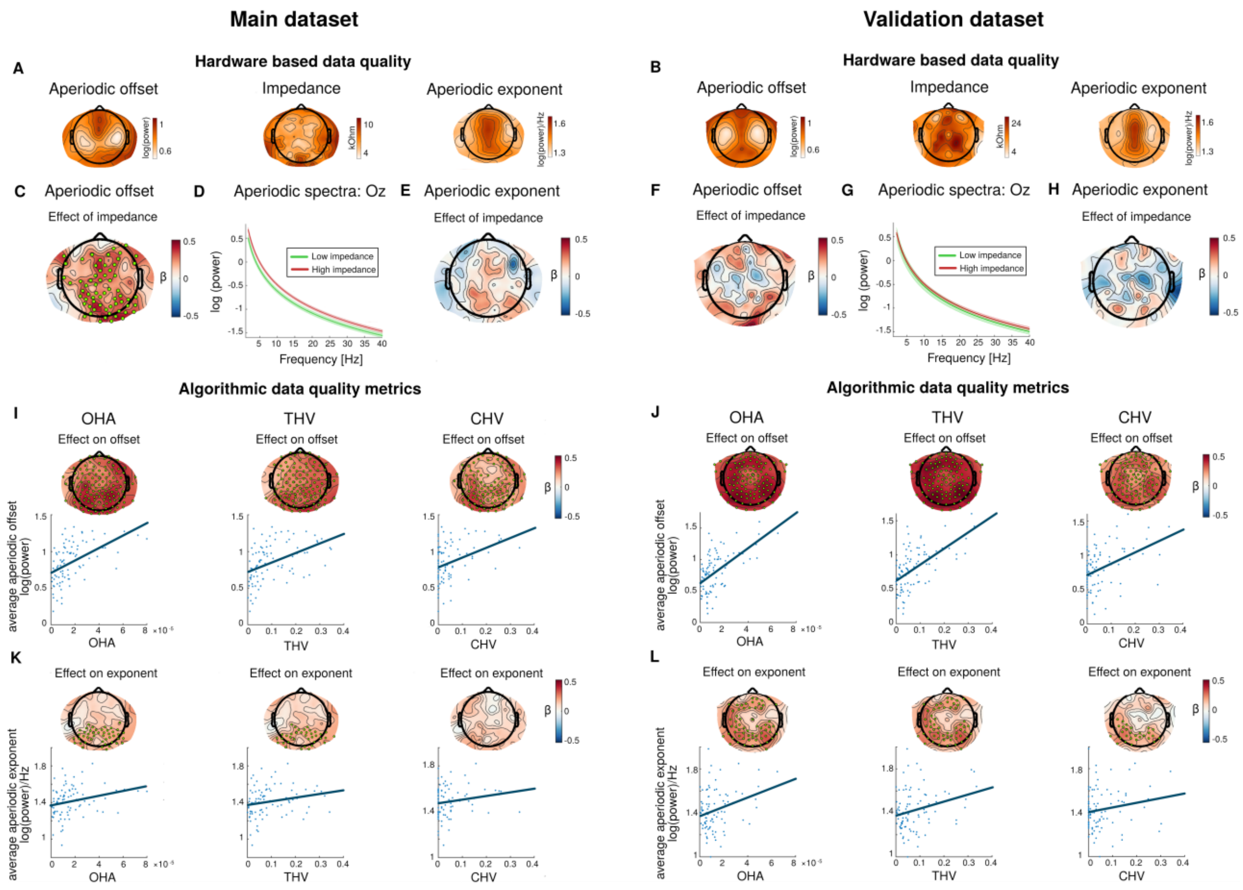


*Supplementary Figure 3: Side by side visualization of data quality impact on the aperiodic parameters in both datasets. Panels A-H illustrate effects of electrode impedances on the aperiodic offset and exponent. Panels A&B show the topographical distribution of the aperiodic offset, exponent, and the impedances of each dataset. Panels C, E, F&H display standardized β's from linear regressions predicting the aperiodic parameters from electrode impedances, green dots mark significant electrode clusters by the CBPT. Panel D&G present median-split aperiodic spectra at electrode Oz, shaded areas indicate standard error of the mean. Panels I-L depict the effects of algorithmic quality metrics (OHA, THV, CHV) on the aperiodic parameters. Topographies show standardized β coefficients, with green dots indicating significant electrodes of the CBPTs. Scatterplots illustrate the relationship between quality metrics and aperiodic parameters averaged across all electrodes.*

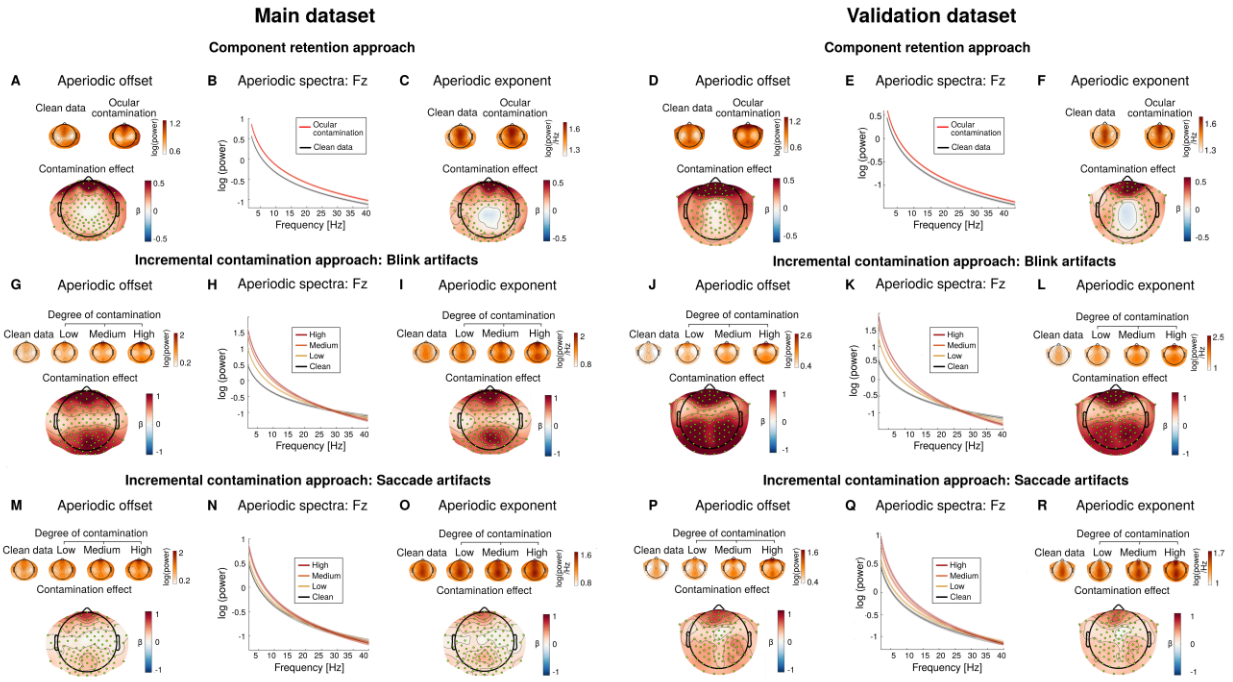


*Supplementary Figure 4: Side by side visualization of ocular artifact impact on the aperiodic parameters in both datasets. Panels A-F illustrate findings from the component retention approach. Small topographies on top of A, C, D&F display aperiodic parameters derived from fully cleaned data (all artifactual independent components (ICs) removed) and partially cleaned (ocular ICs retained) data. Larger maps display standardized regression coefficients from linear mixed-effects models predicting aperiodic signal parameters from ocular contamination; green dots mark electrodes significant in CBPT. Panel B&E show average aperiodic spectra for fully cleaned versus ocular contaminated data, with shaded areas indicating standard error of the mean. Panels G-R present results from the incremental contamination approach for blinks and saccades, using the same visualization schemes as A-F. Contamination levels reflect the proportion of artifactual segments mixed into clean data (clean: 0/15, low: 1/15, medium: 3/15, high: 5/15 segments), with the contamination effect topographies displaying standardized regression coefficients for the clean vs. medium contrast.*

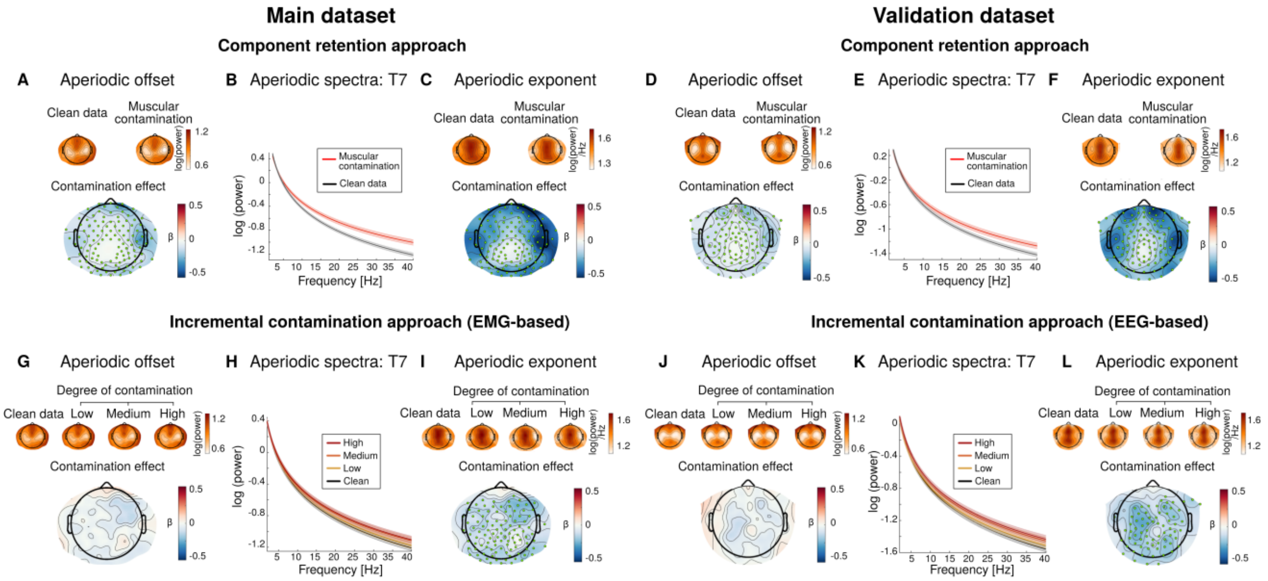


*Supplementary Figure 5: Side by side visualization of muscular artifact influences on the aperiodic parameters in both datasets. Panels A-F illustrate findings from the component retention approach. Small topographies on top of A, C, D&F display aperiodic parameters derived from fully cleaned data (all artifactual ICs removed) and partially cleaned data (muscular ICs retained). Larger maps display standardized regression coefficients from linear mixed-effects models predicting aperiodic signal parameters from muscular contamination; green dots mark electrodes significant in CBPT. Panel B&E show average aperiodic spectra for fully cleaned versus muscular contaminated data, with shaded areas indicating standard error of the mean. Panel G-L present results from the incremental contamination approach for muscular contamination, using the same visualization schemes as A-F. Contamination levels reflect the proportion of artifactual segments mixed into clean data (clean: 0/15, low: 1/15, medium: 3/15, high: 5/15 segments), with the contamination effect topographies displaying standardized regression coefficients for the clean vs. medium contrast.*

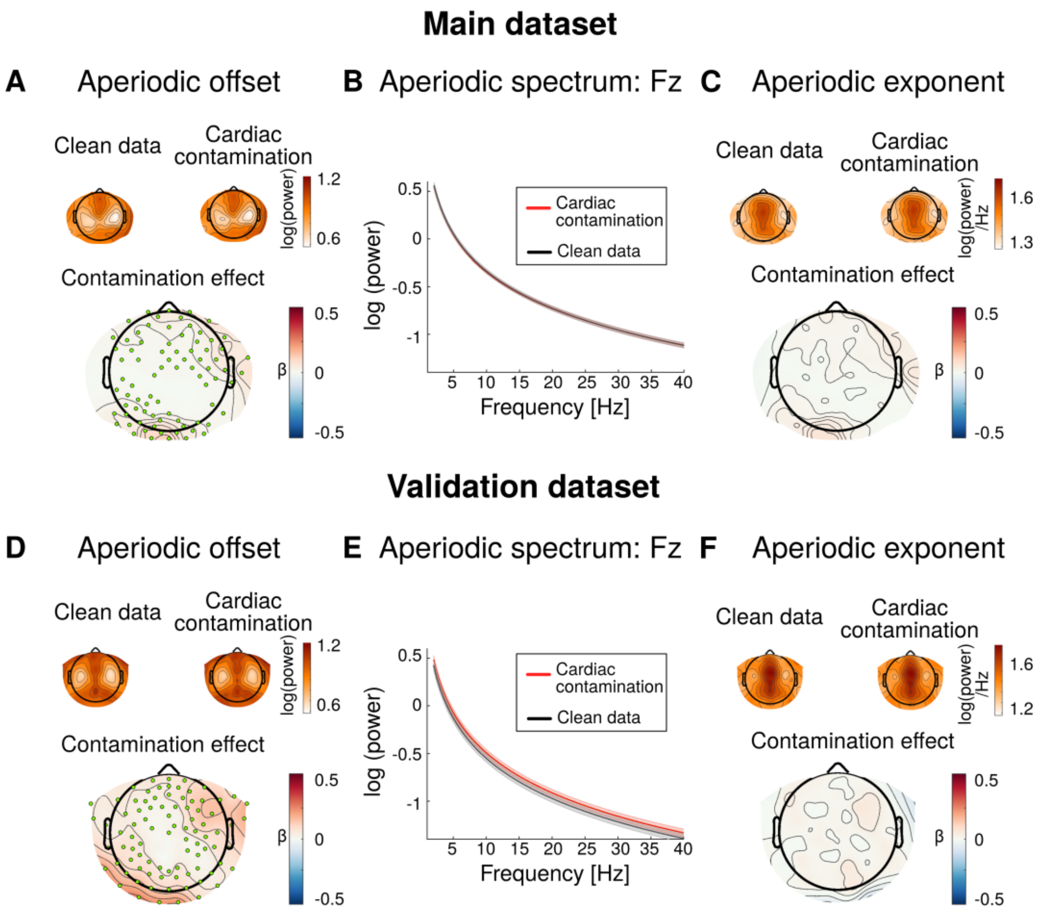


*Supplementary Figure 6: Side by side visualization of cardiac artifact influences on the aperiodic parameters in both datasets from the component retention approach. Note that small topographies on top of A, C, D&F display aperiodic parameters derived from fully cleaned data (all artifactual ICs removed) and partially cleaned (cardiac ICs retained) data. Larger maps display standardized regression coefficients from linear mixed-effects models predicting aperiodic signal parameters from cardiac contamination; green dots mark electrodes significant in CBPT. Panel B&E show average aperiodic spectra for fully cleaned versus cardiac contaminated data, with shaded areas indicating standard error of the mean.*

#### S4. Component retention approach: Ocular artifact removal in eyes-open data

Applying the component retention approach to eyes-open resting-state data yielded highly significant results of ocular contamination on the aperiodic offset, with large standardized beta estimates, averaged across electrodes within the largest significant cluster (main dataset: avg $\beta = 0.73$, $p < 0.001$; validation dataset: avg $\beta = 0.97$, $p < 0.001$). Similar effects were observed on the aperiodic exponent (main dataset: avg $\beta = 0.72$, $p < 0.001$; validation dataset: avg $\beta = 0.90$, $p < 0.001$). Supplementary Figure 7 visualizes these results for both datasets.

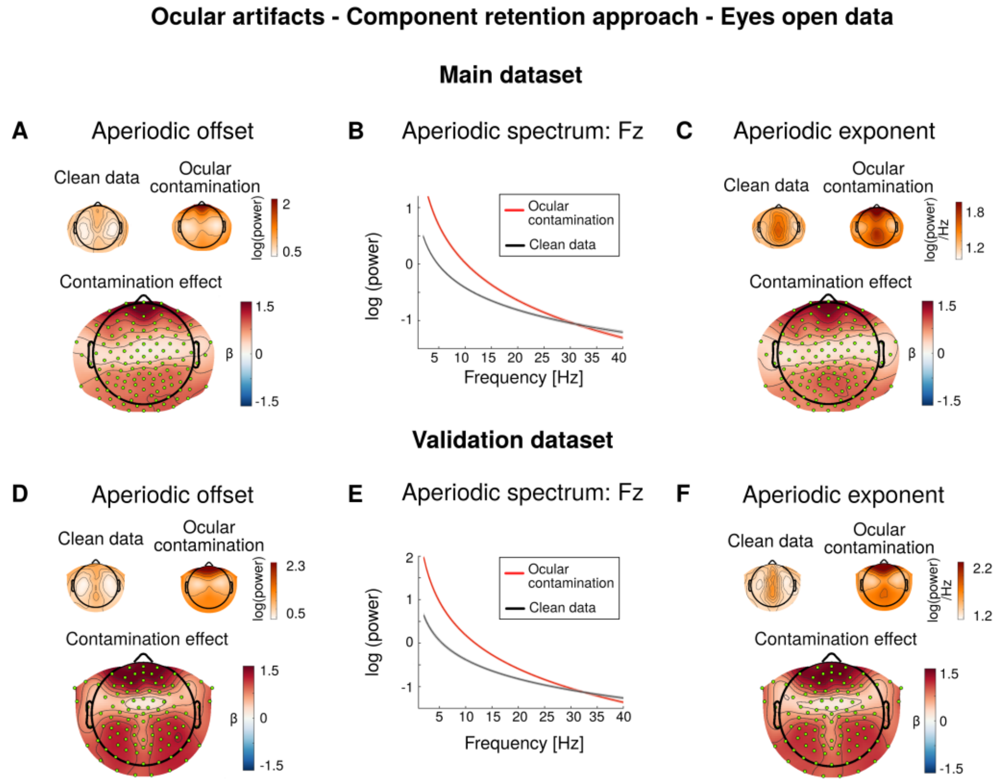


*Supplementary Figure 7: Component retention approach on ocular artifacts, applied to eyes-open resting-state data. Small topographies on top of A&C display aperiodic parameters derived from fully cleaned data (all artifactual ICs removed) and contaminated (ocular ICs retained) data. Larger maps display standardized regression coefficients from linear mixed-effects models predicting aperiodic signal parameters from ocular contamination; green dots mark electrodes significant in CBPT. Panel B shows mean aperiodic spectra for fully cleaned versus ocular contaminated data, with shaded areas indicating standard error of the mean. Panels D-F visualize the results for the validation dataset accordingly.*

**S5. Incremental contamination approach applied to fully cleaned data**

As expected, applying the incremental contamination approach to the fully cleaned data (i.e., after removing all artifact-related ICs) yielded reduced effect sizes compared to those reported in the main manuscript, where ICs of the target artifact category were retained. Importantly, all effects remained statistically significant and of non-negligible magnitude, indicating that standard ICA-based artifact correction does not fully eliminate the influence of physiological artifacts on aperiodic parameter estimation.

For blinks, moderate effects were observed for the aperiodic offset when averaging across the electrodes of the largest significant cluster, with the standardized regression coefficient ($\beta$) reflecting the contrast between medium contamination (3 of 15 segments contaminated) and clean segments (main dataset: avg. $\beta = 0.24$, $p < 0.001$; validation dataset: avg. $\beta = 0.20$, $p < 0.001$). Similar effects were observed for the aperiodic exponent (main dataset: $p = 0.005$, avg. $\beta = 0.20$; validation dataset: avg. $\beta = 0.19$, $p = 0.003$).

The saccade-related results followed the same pattern, showing smaller effects on the aperiodic offset (main dataset: avg. $\beta = 0.08$, $p < 0.001$; validation dataset: avg. $\beta = 0.09$, $p < 0.001$) and on the aperiodic exponent (main dataset: avg. $\beta = 0.09$, $p = 0.017$; validation dataset: avg. $\beta = 0.09$, $p = 0.019$).

For muscular artifacts, consistent with the main manuscript results, no significant effects of muscular contamination were observed on the aperiodic offset. Effects on the aperiodic exponent closely mirrored those reported in the main manuscript: after removal of all muscular ICs, significant effects persisted in both datasets (main dataset, EMG-based detection: $p < 0.001$, avg. $\beta = -0.14$; main dataset, EEG-based detection: avg. $\beta = -0.20$, $p < 0.001$; validation dataset, EEG-based detection: avg. $\beta = -0.17$, $p < 0.001$).

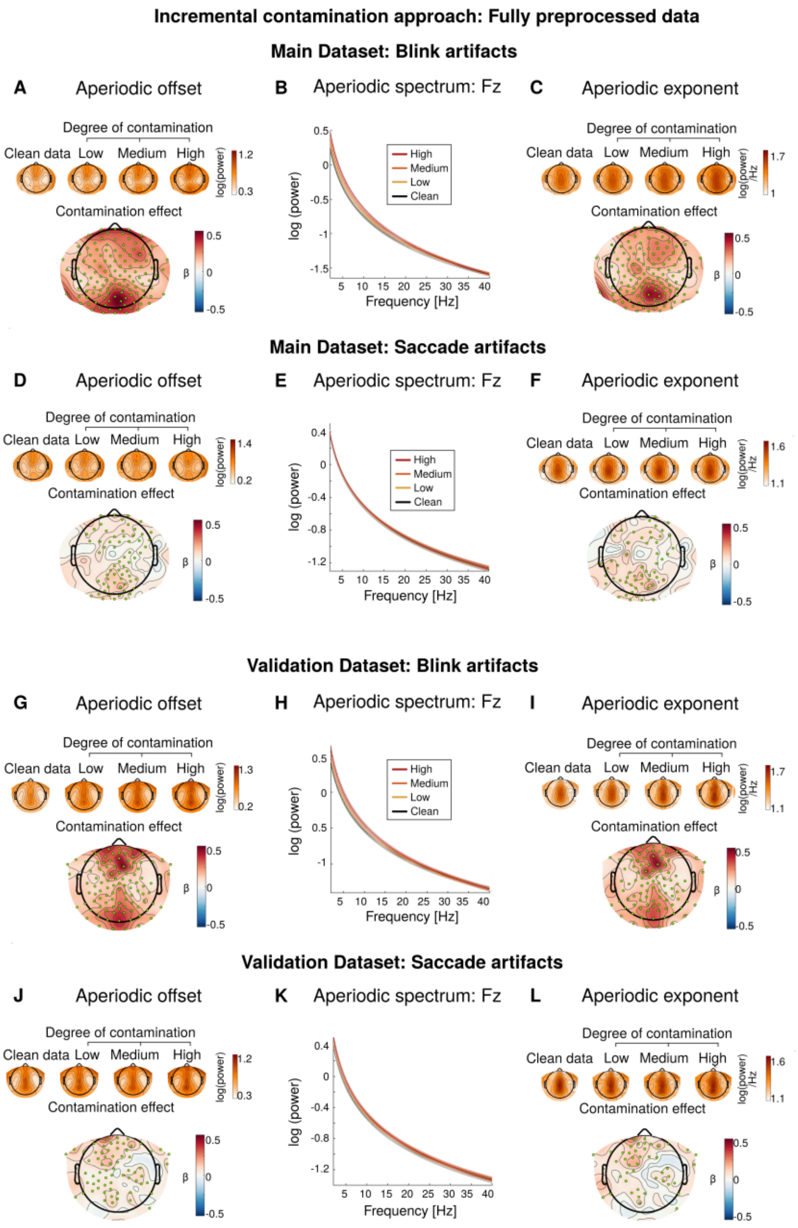


*Supplementary Figure 8: Impact of ocular artifacts on aperiodic parameters based on the incremental contamination approach applied to fully cleaned data (i.e. after removal of all ICs classified as ocular activity). Left column (Panels A/D/G/J) visualizes standardized beta estimates predicting the aperiodic offset by blink and saccade contamination across the scalp, green electrodes highlight significant electrodes in the CBPTs. Here, the contamination effect is visualized by the contrast "clean data" (0 artifactual segments) vs. "medium contaminated data" (3 out of 15 artifactual segments). The right column (Panels C/F/I/L) visualizes the effects on the aperiodic exponent accordingly. The middle column (Panels B/E/H/K) shows average aperiodic spectra for clean or low/medium/high contaminated data (1/3/5 artifactual segments out of 15 total). Shaded error bars represent standard errors of the mean.*

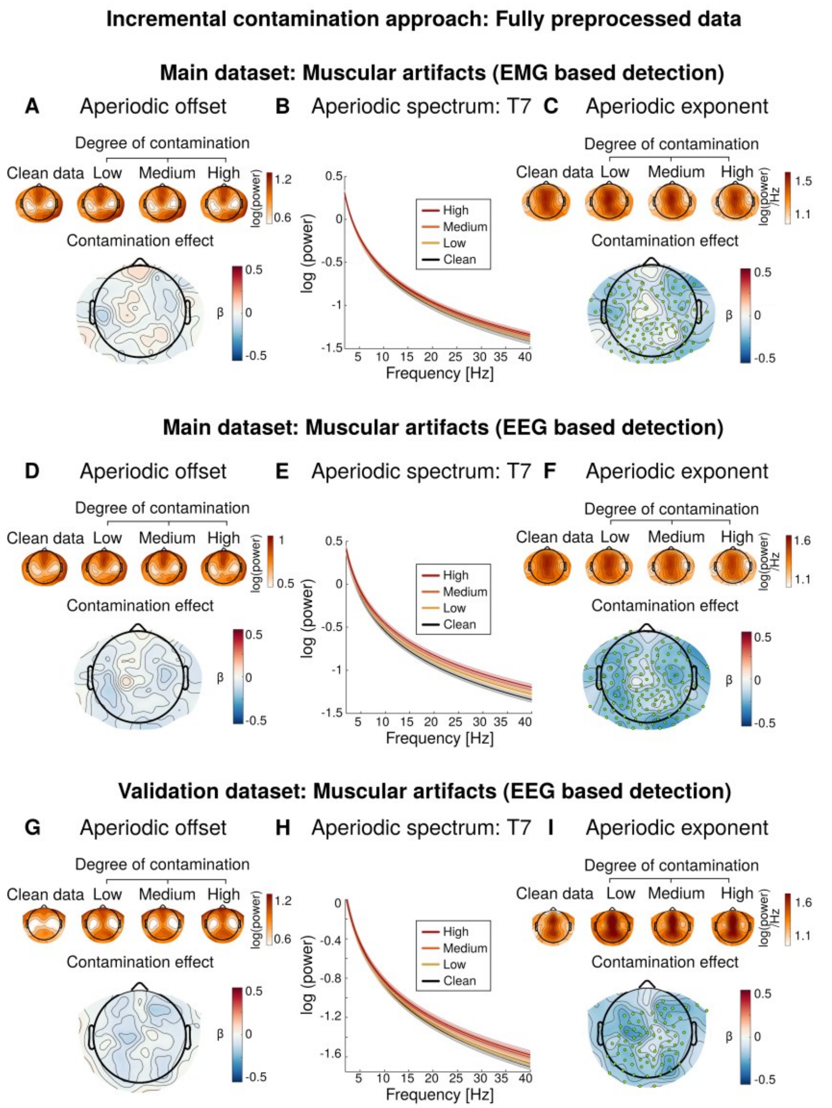


*Supplementary Figure 9: Impact of muscular artifacts on aperiodic parameters based on the incremental contamination approach applied to fully cleaned data (i.e. after removal of all ICs classified as muscular activity). Left column (Panels A/D/G) visualizes standardized beta estimates predicting the aperiodic offset by muscular contamination across the scalp, green electrodes highlight significant electrodes in the CBPTs. Here, the contamination effect is visualized by the contrast "clean data" (0 artifactual segments) vs. "medium contaminated data" (3 out of 15 artifactual segments). For the main dataset, muscular activity is identified either through EMG electrodes, or through an established algorithm based on EEG data (see Methods). The latter is also applied to the validation dataset, which lacks auxiliary EMG electrodes. The right column (Panels C/F/I) visualizes the effects on the aperiodic exponent accordingly. The middle column (Panels B/E/H) shows average aperiodic spectra for clean or low/medium/high contaminated data (1/3/5 artifactual segments out of 15 total). Shaded error bars represent standard errors of the mean.*

**S6. Incremental contamination approach: EEG-based muscle detection in main dataset**

Supplementary Figure 10 visualizes the results of the incremental contamination approach applied to muscular artifact contamination, employing EEG-based artifact classification in the main dataset. Note that the statistical results are reported in the main manuscript (Table 2).

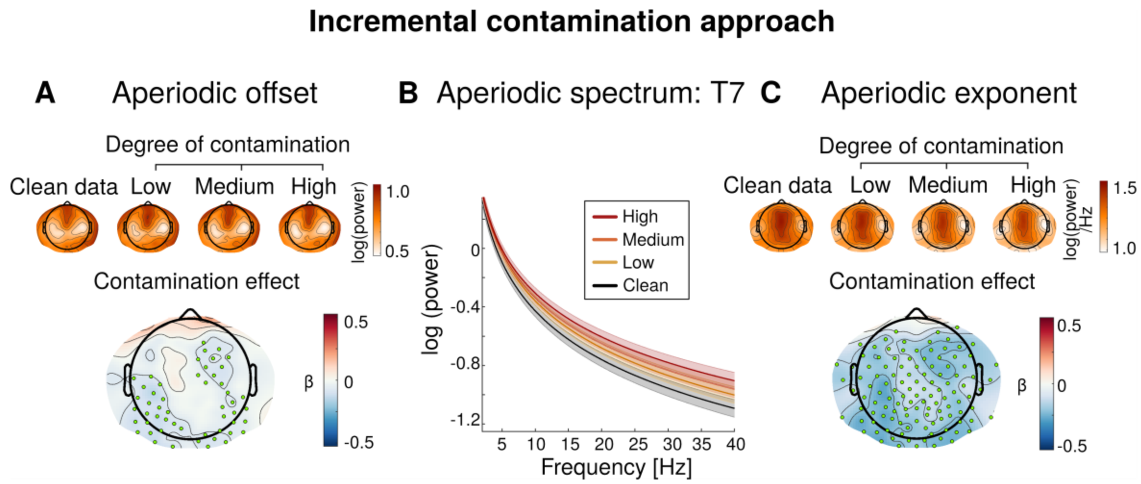


*Supplementary Figure 10: Impact of incremental muscular artifact contamination on the aperiodic parameters, with EEG-based muscular artifact detection in the main dataset. Small topographies in A and C display aperiodic parameters derived from clean or partially contaminated data. Larger maps show standardized regression coefficients from the linear mixed-effects models predicting the aperiodic parameters from muscular contamination; green dots indicate electrodes significant in CBPT. The contamination effects are illustrated by contrast clean data (0 artifactual segments) vs. medium contaminated data (3 out of 15 artifactual segments). Panel B shows mean aperiodic spectra for the clean and partially contaminated data (low: 1/15, medium: 3/15, high: 5/15 contaminated segments), with shaded areas indicating the standard errors of the mean.*

#### S7. Detailed preprocessing pipelines

In a first step, error-prone channels were automatically identified and removed using the eeglab plugin clean_rawdata (http://sccn.ucsd.edu/wiki/Plugin_list_process). Channels were flagged for removal if any of the following criteria were met: a correlation coefficient below 0.85 when compared with an estimate derived from neighboring electrodes; line noise levels exceeding those of all other channels by more than 4 standard deviations; or the presence of a flat-line period longer than 5 seconds. The error-prone electrodes were interpolated using a spherical spline interpolation (EEGLAB function eeg_interp.m) at the end of the preprocessing pipeline before the automatic quality assessment of the EEG files (see below). Data were then high-pass filtered (−6 dB cutoff at 0.5 Hz) and processed with Zapline[2] to eliminate 50 Hz line noise artifacts by removing seven power line components. Next, independent component analysis (ICA) was performed on a temporarily high-pass filtered version of the data (−6 dB cutoff at 1 Hz, resulting ICA weights were later applied to the original 0.5 Hz filtered data). All components were evaluated using the pre-trained ICLabel classifier[1] , which assigns probability scores for line noise, channel noise, muscular activity, ocular activity, cardiac artifacts, brain, and other categories. Rather than immediately removing components with high artifact probabilities, we retained this information for later stages of the analysis pipeline, where we either preserved specific artifact components or excluded all artifactual components (see Figure 1 for an overview). For the subsequent automated quality assessment, however, components with artifact probabilities exceeding 0.8 for line noise, channel noise, muscular activity, ocular activity, or cardiac artifacts were temporarily removed. This temporary removal ensured that quality ratings were based on fully preprocessed data and not confounded by major artifacts. Each EEG file was then evaluated according to four criteria: a file was marked as bad and excluded from further analysis if (1) more than 20% of the data points had amplitudes exceeding 20 μV, (2) over 35% of time points showed a variance greater than 10 μV across channels, (3) 35% or more of the electrodes exhibited high variance (>10 μV), or (4) the overall ratio of excluded electrodes exceeded 0.3 (30%).

After these preprocessing steps, channels positioned outside the standard scalp montage were removed in the validation dataset: These included 10 electrodes placed on the face (primarily recording EOG activity) and 13 electrodes positioned on the chin and neck, which capture minimal brain activity. This resulted in 105 EEG channels in the validation dataset for subsequent analysis. For the main dataset, no channels were removed as it contained 128 standard scalp EEG channels. Finally, the data were re-referenced to a common average reference.

**S8. Specparam model fits per analysis**

Because each analysis type used a specific subsample of the data (and varying degree of artifact contamination), the model fits of the specparam algorithm[3] were evaluated separately and are presented in Supplementary Table 2. Importantly, also for the incremental contamination analysis, in which the PSD is estimated based on 15 one-second segments, very high model fits were obtained. When further dividing this into the sub-analysis of clean (0/15) / low (1/15) / medium (3/15) / high contamination (5/15), no systematic differences in specparam fits were observed.

**Supplementary Table 2**

*Average model fits of the specparam algorithm for each analysis subtype*

| | Main dataset: Mean specparam fit $R^2$ (sd) | Validation dataset: Mean specparam fit $R^2$ (sd) |
|---|---|---|
| Data quality and component retention approach (all artifact ICs removed) | 0.99 (0.01) | 0.99 (0.02) |
| Component retention approach: Ocular artifacts retained | 0.99 (0.01) | 0.99 (0.02) |
| Component retention approach: Muscular artifacts retained | 0.99 (0.03) | 0.98 (0.03) |
| Component retention approach: Cardiac artifacts retained | 0.99 (0.01) | 0.99 (0.01) |
| Incremental contamination approach: Blink artifacts | 0.96 (0.07) | 0.96 (0.06) |
| Incremental contamination approach: Saccade artifacts | 0.96 (0.07) | 0.96 (0.06) |
| Incremental contamination approach: Muscular artifacts | 0.97 (0.05) | 0.96 (0.06) |

#### S9. Distribution of impedances and algorithmic quality metrics

Supplementary Figure 11 visualizes the distribution of electrode impedances (across all electrodes and participants) and algorithmic quality metrics. Only impedance values within the ranges used for the analyses are included in the plots (Main dataset: 0-20 kΩ, Validation dataset:0-50 kΩ).

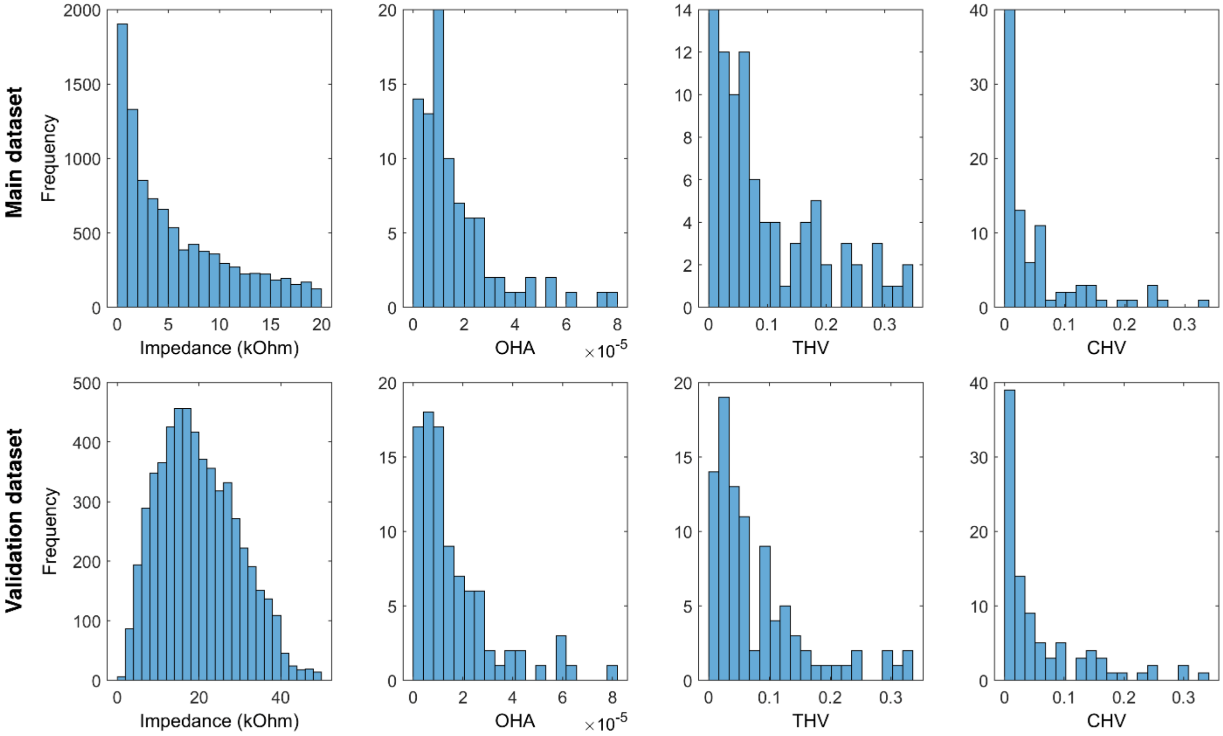


*Supplementary Figure 11: Distribution of electrode impedances and algorithmic quality measures (overall high amplitude, OHA; timepoints of high variance, THV; channels of high variance, CHV) in each dataset.*

#### S10. Relation between algorithmic quality metrics and electrode impedances

The relationships between the investigated quality measures are depicted in Supplementary Table 3. In both datasets, electrode impedances were not significantly related to any of the algorithmic quality metrics. In contrast, the three different algorithmic quality metrics were strongly interconnected.

**Supplementary Table 3**

*Pearson correlations between the algorithmic quality metrics and impedance measures*

| | Impedances: *r* (*p*) | OHA: *r* (*p*) | THV: *r* (*p*) |
|---|---|---|---|
| OHA | 0.08 (0.44) / -0.03 (0.85) | | |
| THV | 0.17 (0.10) / -0.06 (0.66) | 0.70 (<0.001) / 0.97 (<0.001) | |
| CHV | -0.02 (0.86) / -0.03 (0.86) | 0.83 (<0.001) / 0.91 (<0.001) | 0.73 (<0.001) / 0.83 (<0.001) |

*Note: Impedances are averaged across all electrodes. Values are reported as main dataset / validation dataset.*

#### S11. Muscular artifact detection algorithm

The EEG-based muscular artifact detection algorithm was adapted from an established algorithm[4], combined with the automated muscular artifact detection algorithm implemented in MNE-Python[5]. First, PCA was applied to all EEG channels and the first principal component was extracted for further analyses, as this component typically contains high amplitude artifacts[6]. The resulting time series was filtered to 65-115 Hz to further reduce the influence of brain activity on the extracted signal and enhance signal components related to muscular activity. Subsequently, the envelope of this signal was calculated using Hilbert transformation and demeaned within each of the 5 concatenated eyes-closed recording blocks. Instead of directly thresholding this envelope, we applied a low pass filter at 4 Hz to remove highly transient peaks which are unlikely to reflect muscular activity, as proposed in the MNE-Python muscular artifact detection algorithm. Finally, the resulting signal was z-transformed and thresholded at $z > 2$, determined by visual inspection across all participants, to identify periods of muscular activity based on extreme values in the distribution. The visual inspection of the annotations revealed that while this approach reliably detected muscular artifacts in participants with pronounced contamination, in very clean recordings however, segments without visible artifacts were occasionally misclassified as contaminated. To mitigate this issue, an additional constraint was imposed by retaining only those segments where the envelope of the first principal component exceeded 20 µV, thereby ensuring that only clear muscular activity was detected. The data were then segmented to one-second epochs centered around the annotated data. If the length of a continuous annotation was shorter than one-second, one segment was extracted around the center of the annotation period. If an annotation lasted longer than one-second, multiple non-overlapping one-second segments were extracted each containing at least 50% of the annotated period, thereby avoiding segments near annotation boundaries with minimal contamination.

For the EMG-based muscular artifact detection, we first calculated the envelope of each of the three EMG channels using Hilbert transformation and subsequently applied a low pass filter of 4 Hz to this envelope to remove transient noisy peaks. The resulting signals were z-transformed and thresholded at $z > 5$ (determined by visual inspection, analogous to the EEG-based approach), segments were annotated as contaminated if activity in at least one of the three EMG channels exceeded this threshold. Finally, the same segmentation around these annotation periods was applied as described above. Because the frontal EMG channels were also sensitive to artifacts from eye movements and blinks, we applied the same annotation procedure also to the HEOG and VEOG channels. If any segment marked as muscular activity was also annotated as high EOG activity (either in the HEOG or the VEOG channel), the segment was not included for further analyses.

**S12. Robustness analysis of the statistical correction approach**

To ensure that the simulation results reported in the main manuscript were not dependent on a specific random split, we repeated the analysis across 5,000 permutations, each using a different random seed for participant assignment and segment sampling. For each permutation, we extracted the standardized beta estimate and the p value for the group comparison, both before and after regressing out the number of artifact-contaminated segments.

As expected, groups with matched contamination levels (Group 1 vs. Group 2 Low) showed no systematic difference in aperiodic exponents (G1: mean = 1.521, sd = 0.024, G2 Low: mean = 1.521, sd = 0.024), but the group with different commination levels did (G2 higher: mean = 1.60, sd = 0.025, see Supplementary Figure 12A, top). After regressing out artifacts, all groups showed equal aperiodic exponents (G1: mean = 1.546, sd = 0.021, G2 Low: mean = 1.546, sd = 0.029. G2 higher: mean = 1.547, sd = 0.021, Supplementary Figure 12A, bottom). This confirmed that differential artifact contamination reliably induces spurious group differences that can be effectively mitigated through statistical correction, independent of the random splits. Supplementary Figure 12B further visualizes the statistical results of each permutation, comparing aperiodic exponents of Group1 (low contamination) and Group 2 (higher contamination) before and after regressing out the number of blink artifacts, employing the same linear models as reported in the main manuscript.

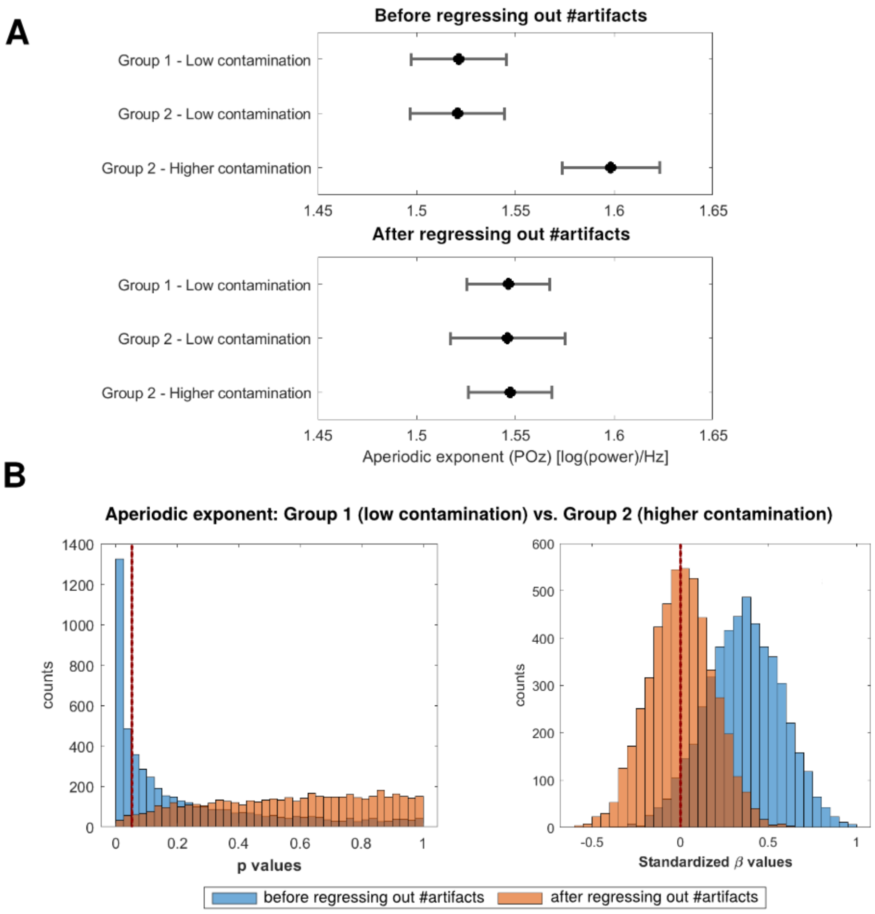


Supplementary Figure 12: Robustness of the statistical correction approach across 5,000 permutations. Panel A displays mean aperiodic exponents (± sd) at electrode POz for each group, averaged across all permutations, before (top) and after (bottom) regressing out the number of blink artifacts. Panel B shows the distributions of p values (left, red dashed line indicates $p$=0.05) and standardized β estimates (right, red dashed line indicates $\beta=0$) from the comparison of Group 1 (low contamination) and Group 2B (higher contamination) across all permutations, before (orange) and after (blue) statistical correction.
